# World Influence of Infectious Diseases from Wikipedia Network Analysis

**DOI:** 10.1101/424465

**Authors:** Guillaume Rollin, José Lages, Dima L. Shepelyansky

**Affiliations:** Institut UTINAM, Observatoire des Sciences de l’Univers THETA, CNRS, Université de Bourgogne Franche-Comté, 25030 Besançon, France; Laboratoire de Physique Théorique, IRSAMC, Université de Toulouse, CNRS, UPS, 31062 Toulouse, France

## Abstract

We consider the network of 5 416 537 articles of English Wikipedia extracted in 2017. Using the recent reduced Google matrix (REGOMAX) method we construct the reduced network of 230 articles (nodes) of infectious diseases and 195 articles of world countries. This method generates the reduced directed network between all 425 nodes taking into account all direct and indirect links with pathways via the huge global network. PageRank and CheiRank algorithms are used to determine the most influential diseases with the top PageRank diseases being Tuberculosis, HIV/AIDS and Malaria. From the reduced Google matrix we determine the sensitivity of world countries to specific diseases integrating their influence over all their history including the times of ancient Egyptian mummies. The obtained results are compared with the World Health Organization (WHO) data demonstrating that the Wikipedia network analysis provides reliable results with up to about 80 percent overlap between WHO and REGOMAX analyses.

## Introduction

*Infectious diseases account for about 1 in 4 deaths worldwide*, *including approximately two-thirds of all deaths among children younger than age 5* (***NIH, 2018***). Thus the understanding of the world influence of infectious diseases is an important challenge. Here we apply the mathematical statistical methods originated from computer and network sciences using the PageRank and other Google matrix algorithms developed at the early stage of search engines development (***Brin and Page, 1998*; *Langville and Meyer, 2012***). These methods are applied to English Wikipedia edition which is considered as a directed network generated by hyperlinks (citations) between articles (nodes). Nowadays, the free online encyclopedia supersedes old ones such as ***Encyclopaedia Britannica (2018)*** in volume and in quality of articles devoted to scientific topics (***Giles, 2005***). For instance, Wikipedia articles devoted to biomolecules are actively maintained by scholars of the domain (***Butler, 2008*; *Callaway, 2010***). The academic analysis of information contained by Wikipedia finds more and more applications as reviewed in ***Reagle Jr. (2012)*; *Nielsen (2012)*.**

The Google matrix analysis, associated to the PageRank algorithm, initially invented by Brin and Page to efficiently rank pages of the World Wide Web (***Brin and Page, 1998***), allows to probe the network of Wikipedia articles in order to measure the influence of every articles. The efficiency of this approach for Wikipedia networks has been demonstrated by ranking historical figures on a scale of 35 centuries of human history and by ranking world universities (***Zhirov et al., 2010*; *Eom et al., 2015*; *Lages et al., 2016*; *Katz and Rokach, 2017*; *Coquidé et al., 2018***). This approach produced also reliable results for the world trade during last 50 years reported by the UN COMTRADE database and other directed networks (***Ermann et al., 2015***).

Recently, the reduced Google matrix method (REGOMAX) has been proposed using parallels with quantum scattering in nuclear physics, mesoscopic physics, and quantum chaos (***Frahm and Shepelyansky, 2016*; *Frahm et al., 2016***). This method allows to infer hidden interactions between a set of *n_r_* nodes selected from a huge network taking into account all indirect pathways between these *n_r_* nodes via the huge remaining part of the network. The efficiency of REGOMAX has been demonstrated for analysis of world terror networks and geopolitical relations between countries from Wikipedia networks (***El Zant et al., 2018a***,b). The efficient applications of this approach to the global biological molecular networks and their signaling pathways are demonstrated in (***Lages et al., 2018***).

In this work we use REGOMAX method to investigate the world influence and importance of infectious diseases constructing the reduced Google matrix from English Wikipedia network with all infectious diseases and world countries listed there.

The paper is constructed as follows: the data sets and methods are described in Section II, Results are presented in Section III and discussion is given in Section IV; Appendix contains Tables 1, 2, 3, 4, 5; additional data are presented at ***Wiki4InfectiousDiseases (2018)***.

## Description of data sets and methods

### English Wikipedia Edition network

We consider the English language edition of Wikipedia collected in May 2017 (***Frahm and Shepelyansky, 2017***) containing *N* = 5 416 537 articles (nodes) connected through *n_l_* = 122 232 932 hyperlinks between articles. From this data set we extract the *n_d_* = 230 articles devoted to infectious diseases (see Tab. 1, Tab. 2) and the *n_c_* = 195 articles devoted to countries (sovereign states, see Tab. 3). The list of infectious diseases is taken from ***Wikipedia (2018d)*** and the list of sovereign states of 2017 is taken from ***Wikipedia (2017)***. Thus the size of the reduced Google matrix is *n_r_* = *n_d_* + *n_c_* = 425. This subset of *n_r_* articles is embedded in the global Wikipedia network with *N* nodes. All data sets are available at ***Wiki4InfectiousDiseases (2018)***.

### Google matrix construction

The construction of Google matrix *G* is described in detail in ***Brin and Page (1998)***; ***Langville and Meyer (2012***); ***Ermann et al. (2015)***. In short, the Google matrix *G* is constructed from the adjacency matrix *A_ij_* with elements 1 if article (node) *j* points to article (node) *i* and zero otherwise. The Google matrix elements take the standard form *G_ij_* = *αS_ij_* + (1 – *α*)/*N* (***Brin and Page, 1998*; *Langville and Meyer, 2012***; ***Ermann et al., 2015***), where *S* is the matrix of Markov transitions with elements *S_ij_* = *A_ij_* /*k_out_*(*j*). Here *k_out_*(*j*) = 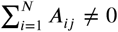 is the out-degree of node *j* (number of outgoing links) and *S_ij_* = 1/*N* if *j* has no outgoing links (dangling node). The parameter 0 < *α* < 1 is the damping factor. For a random surfer, jumping from one node to another, it determines the probability (1 – *α*) to jump to any node; below we use the standard value *α* = 0.85 (***Langville and Meyer, 2012***).

The right eigenvector of *G* satisfies the equation *GP* = *λP* with the unit eigenvalue *λ* = 1. It gives the PageRank probabilities *P (j*) to 1nd a random surfer on a node *j* and has positive elements Σ*_j_ P (j*) = 1). All nodes can be ordered by decreasing probability *P* numbered by PageRank index *K* = 1, 2, … *N* with a maximal probability at *K* = 1 and minimal at *K* = *N*. The numerical computation of *P (j*) is efficiently done with the PageRank algorithm described in ***Brin and Page (1998)***; ***Langville and Meyer (2012)***.

It is also useful to consider the network with inverted direction of links. After inversion the Google matrix *G*^∗^ is constructed within the same procedure with *G*^∗^*P* ^∗^ = *P* ^∗^. This matrix has its own PageRank vector *P* ^∗^(*j*) called CheiRank (***Chepelianskii, 2010***) (see also ***Zhirov et al. (2010)*; *Ermann et al. (2015)***). Its probability values can be again ordered in a decreasing order with CheiRank index *K*^∗^ with highest *P* ^∗^ at *K*^∗^ = 1 and smallest at *K*^∗^ = *N*. On average, the high values of *P (P* ^∗^) correspond to nodes with many ingoing (outgoing) links (***Ermann et al., 2015***).

### Reduced Google matrix analysis

Reduced Google matrix is on-structed for a selected subset of nodes (articles) following the method described in ***Frahm and Shepelyansky (2016)*; *Frahm et al. (2016)*; *Lages et al. (2018)***. It is based on concepts of scattering theory used in different 1elds including mesoscopic and nuclear physics, and quantum chaos (see Refs. in ***Frahm and Shepelyansky (2016)***). It captures in a *n_r_*-by-*n_r_* Perron-Frobenius matrix the full contribution of direct and indirect interactions happening in the full Google matrix between the *n_r_* nodes of interest. Also the PageRank probabilities of selected *n_r_* nodes are the same as for the global network with *N* nodes, up to a constant multiplicative factor taking into account that the sum of PageRank probabilities over *n_r_* nodes is unity. The elements of reduced matrix *G*_R_(*i, j*) can be interpreted as the probability for a random surfer starting at web-page *j* to arrive in web-page *i* using direct and indirect interactions. Indirect interactions refer to paths composed in part of web-pages different from the *n_r_* ones of interest. The intermediate computation steps of *G*_R_ offer a decomposition of *G*_R_ into matrices that clearly distinguish direct from indirect interactions: *G*_R_ = *G*_rr_ + *G*_pr_ + *G*_qr_ (***Frahm et al., 2016***). Here *G*_rr_ is given by the direct links between selected *n_r_* nodes in the global *G* matrix with *n* nodes. In fact, *G*_pr_ is rather close to the matrix in which each column is given by the PageRank vector *P_r_*, ensuring that PageRank probabilities of *G*_R_ are the same as for *G* (up to a constant multiplier). Thus *G*_pr_ doesn’t provide much information about direct and indirect links between selected nodes. The component playing an interesting role is *G*_qr_, which takes into account all indirect links between selected nodes appearing due to multiple paths via the global network nodes *N* (see ***Frahm and Shepelyansky (2016)***; ***Frahm et al. (2016)***). The matrix *G*_qr_ = *G*_qrd_ + *G*_qrnd_ has diagonal (*G*_qrd_) and non-diagonal (*G*_qrnd_) parts. Thus *G*_qrnd_ describes indirect interactions between nodes. The explicit formulas as well as the mathematical and numerical computation methods of all three components of *G*_R_ are given in ***Frahm and Shepelyansky (2016)***; ***Frahm et al. (2016)***; ***Lages et al. (2018)***.

After obtaining the matrix *G*_R_ and its components we can analyze the PageRank sensitivity in respect to specific links between *n_r_* nodes. To measure the sensitivity of a country *c* to a disease *d* we change the matrix element *G*_R_(*d* → *c*) by a factor (1 + *δ*) with *δ* ≪ 1, we renormalize to unity the sum of the column elements associated with disease *d*, and we compute the logarithmic derivative of PageRank probability *P (c*) associated to country *c*: *D*(*d* → *c, c*) = *d* ln *P (c*)/*dδ* (diagonal sensitivity). It is also possible to consider the nondiagonal (or indirect) sensitivity *D*(*d* → *c, c*’) = *d* ln *P (c*’)/*dδ* when the variation is done for the link from *d* to *c* and the derivative of PageRank probability is computed for another country *c*’. This approach was already used in ***El Zant et al. (2018a,b)*** showing its efficiency.

## Results

### Network of direct links

For the reduced Google matrix analysis we have *n_r_* = 425 selected nodes of countries (195) and infectious diseases (230). The diseases are attributed to 7 groups corresponding to the standard disease types as it is given in Tab. 1, Tab. 2. These *n_r_* nodes constitute a subnetwork embedded in the huge global English Wikipedia network with more than 5 million nodes. This subnetwork is shown in Fig. 1 which has been generated with Cytoscape software (***Shannon et al., 2003***). In Fig. 1 black arrow links represent the nonzero elements of adjacency matrix between the selected *n_r_* nodes. The image of this adjacency matrix is shown in Fig. 2 where white pixels depicted a link between two nodes. In this picture, nodes are ordered with respect to the PageRank order in each subgroup: American countries, European countries, Asian countries, African countries, Oceanian countries, bacterial diseases, viral disease, parasitic diseases, fungal diseases, multiple origins diseases, prionic diseases and other kind of disease origins. There are visibly more links inside subgroups but links between groups are also significant. Fig. 1 gives us the global view of network of direct links, shown in Fig. 2, corresponding to the component *G*_rr_ of the reduced Google matrix. We see that countries are located in the central part of the network of Fig. 1 since they have many ingoing links. While it is useful to have such a global view it is clear that it does not take into account the indirect links appearing between *n_r_* nodes due to pathways via the complementary network part with a huge number of nodes *N* – *n_r_* ≃ *N*. The indirect links emerging between *n_r_* from this indirect pathways are analyzed in the frame of REGOMAX method below.

**Figure 1.**
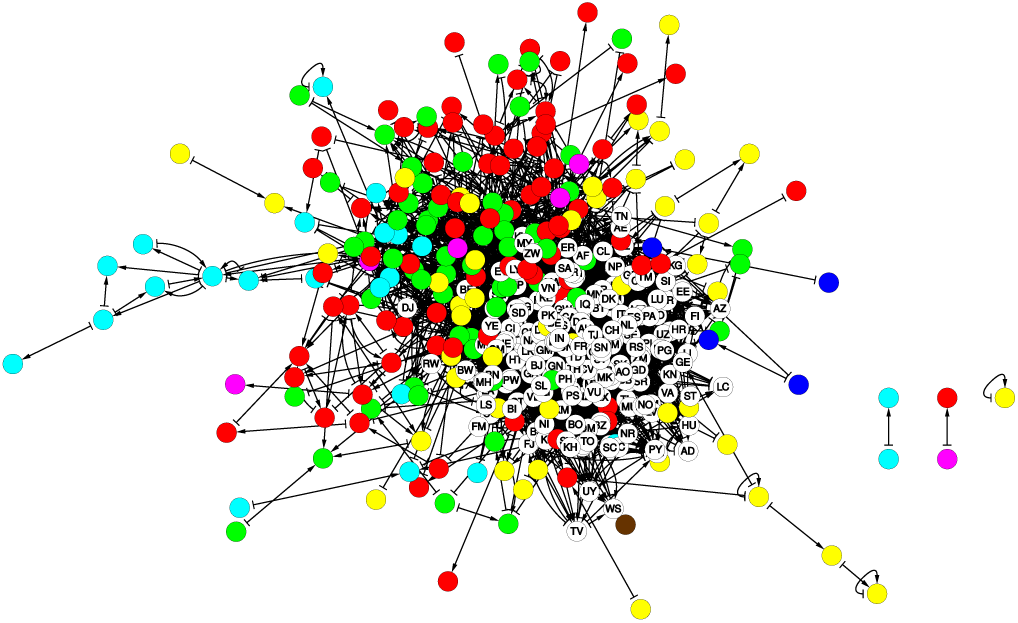
Subnetwork of the 425 articles devoted to countries and infectious diseases in 2017 English Wikipedia. The bulk of the other Wikipedia articles is not shown. Articles devoted to countries are presented by empty nodes with country codes (see Tab. 3). Articles devoted to infectious diseases are presented by colored nodes with the following color code: bacterial diseases, viral diseases, parasitic diseases, fungal diseases, prionic diseases, diseases with multiple origins, and other kind of diseases (see Tab. 1. Tab. 2). Network drawn with Cytoscape (***Shannon et al., 2003***).

**Figure 2.**
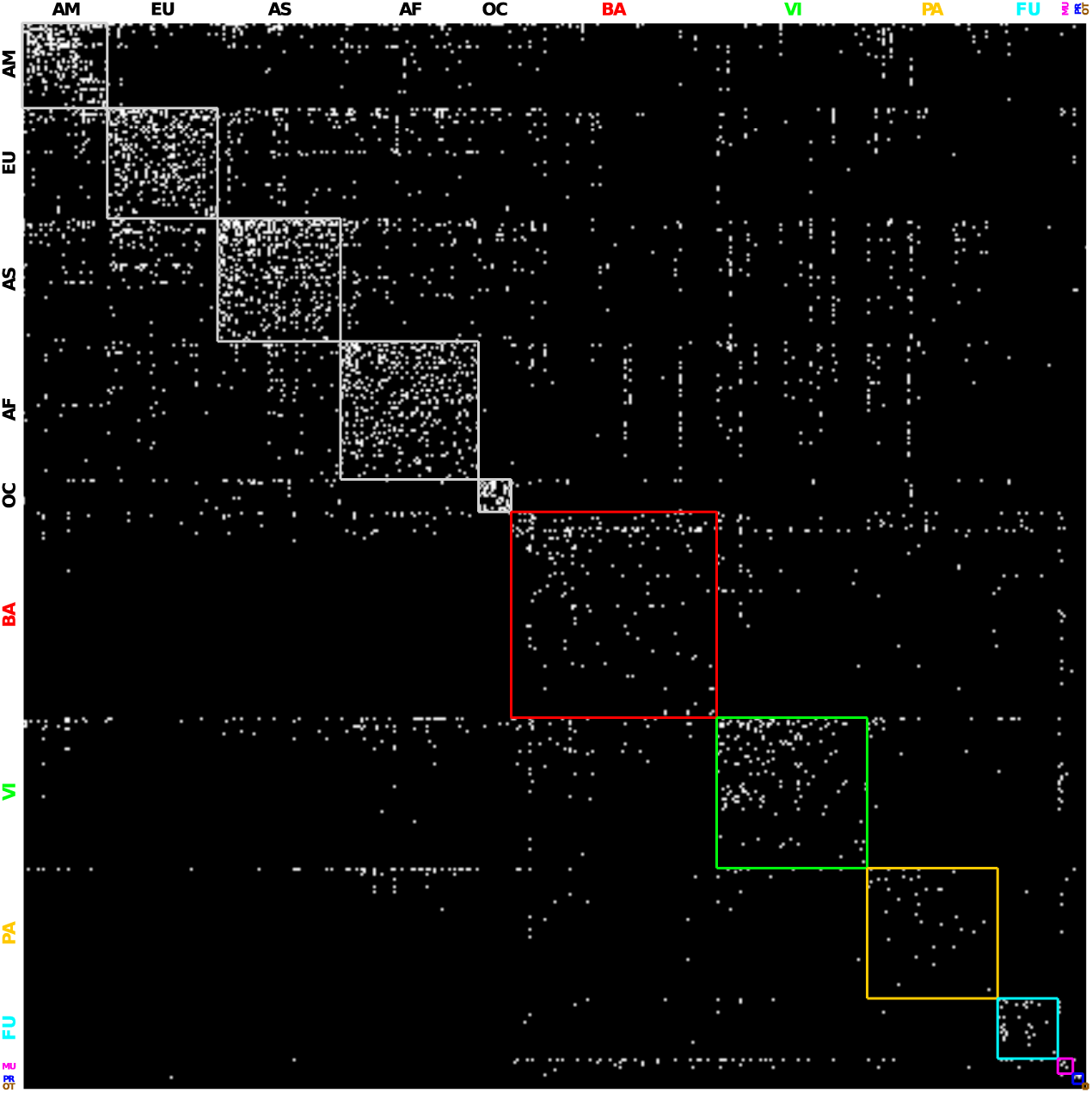
Adjacency matrix of the subnetwork of the 425 articles devoted to countries and infectious diseases in 2017 English Wikipedia. White (black) pixels represent existing (absent) links. In horizontal and vertical axis the articles are ordered by continent for countries (AM for Americas, EU for Europe, AS for Asia, AF for Africa, and OC for Oceania) and by disease type for diseases (**BA** for bacterial, **VI** for viral, **PA** for parasitic, **FU** for fungal, **MU** for multiple type, **PR** for prionic, and **OT** for other type). Inside each block of continent and disease type the articles are ordered according to PageRank algorithm (see Tabs. 1 and 3). From top left to bottom right: block diagonal elements corresponding to intra-continent links are delimited by light gray squares. Also block diagonal elements corresponding to intra-disease type links are delimited by squares with color contour. Off diagonal blocks indicates links between diseases of different type, links between countries of different continent, or links between diseases and countries.

### PageRank and CheiRank of the reduced network nodes

At first we compute the PageRank and CheiRank probabilities for the global network with *N* nodes attributing to each node PageRank and CheiRank indexes ***K*** and ***K***^∗^. For selected *n* nodes the results of PageRank are shown in Tab. 2. As usual (see ***Zhirov et al. (2010)*; *Ermann et al. (2015)***), the countries are taking the top PageRank positions with US, France, Germany, etc at ***K*** = 1, 2, 3, etc as shown in Tab. 3. In the list of *n_r_* nodes the infectious diseases start to appear from ***K*** = 106 with Tuberculosis (Tab. 2). If we consider only infectious diseases ordered by their disease PageRank index ***K**_d_* then we obtain at the top Tuberculosis, HIV/AIDS, Malaria, Pneumonia, Smallpox at first positions with ***K**_d_* = 1, 2, 3, 4, 5 (see Tab. 2). It is clear that PageRank order gives at the top positions severe infectious diseases which are (were for Smallpox) very broadly spread worldwide.

In Fig. 3 we show the location of selected *n_r_* nodes on the global (***K, K***^∗^) plane of density of Wikipedia articles (see details of this representation in ***Zhirov et al. (2010)***; ***Ermann et al. (2015)***). Here the positions of countries are shown by white circles and diseases by color circles. The countries are taking the top positions since they have many ingoing links from variety of other articles. The infectious diseases are located on higher values of ***K, K***^∗^ even if some diseases are overlapping with the end list of countries (see Tab. 2).

**Figure 3.**
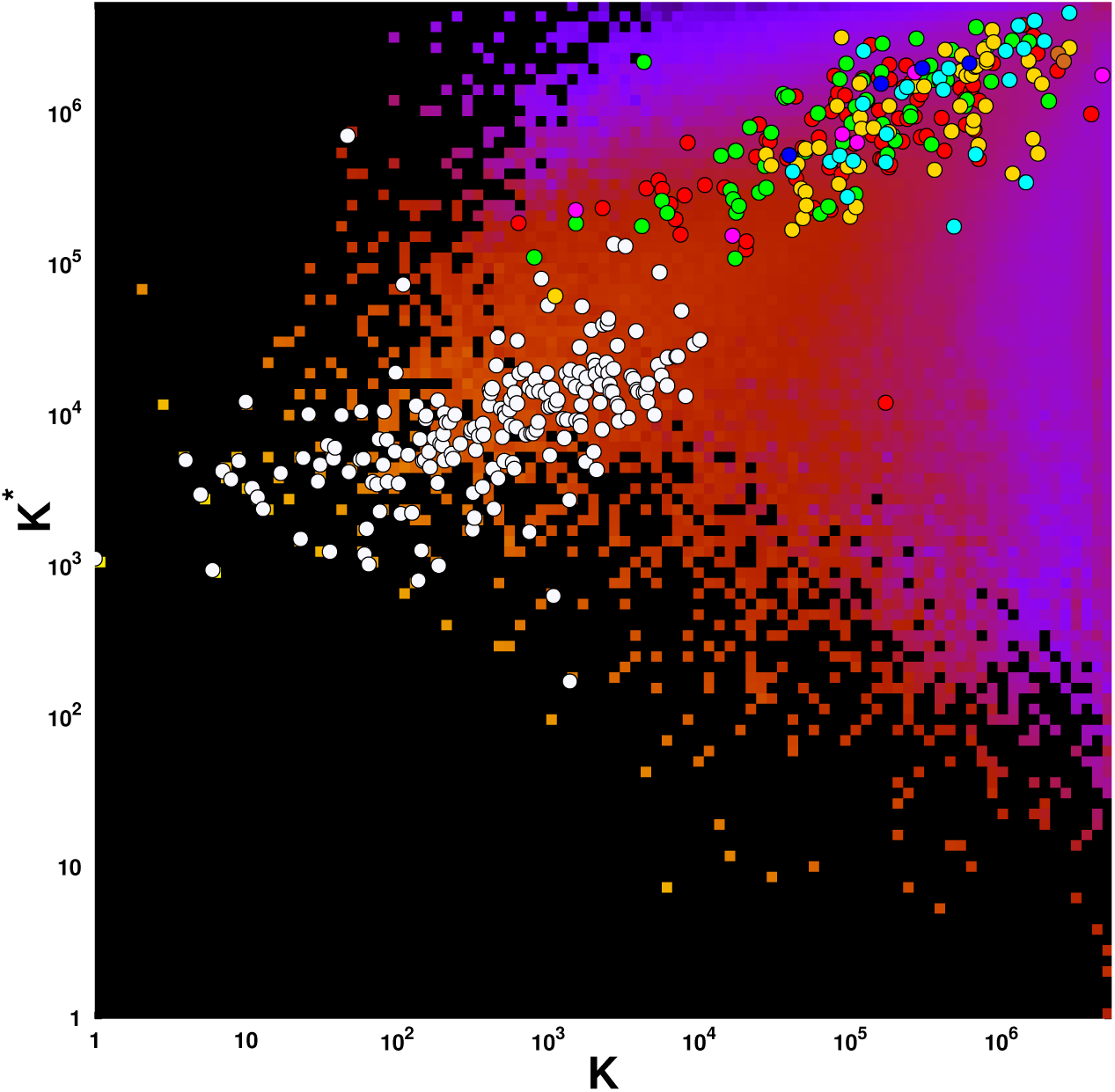
Density of articles of English Wikipedia 2017 on PageRank ***K*** – CheiRank ***K***^∗^ plane. Data are averaged over a 100 × 100 grid spanning the (log_10_ ***K***, log_10_ ***K***^∗^) ∈ [0, log_10_ *N*] × [0, log_10_ *N*] domain. Density of articles ranges from very low density (purple tiles) to very high density (bright yellow tiles). The cells without articles are represented by black tiles. The superimposed white (colored) circles give the positions of countries (infectious diseases) listed in Tab. 3 (Tab. 1). The color code for infectious diseases is the same as in Fig. 1.

All *n_r_* = 425 selected articles can be ordered by their local PageRank and CheiRank indexes ***K**_r_* and ***K**_r_* ^∗^ which range from 1 to *n_r_* = 425. Their distribution in the local PageRank-CheiRank plane is shown in Fig. 4. As discussed previously, countries are at the top ***K**_r_, **K**_r_*^∗^ positions. The names of top PageRank diseases are marked on the figure. The most communicative articles of infectious diseases are those with top ***K**_r_*^∗^ positions. Thus the top CheiRank disease is Burkholderia due to many outgoing links present in this article. The next ones are Malaria and HIV/AIDS.

**Figure 4.**
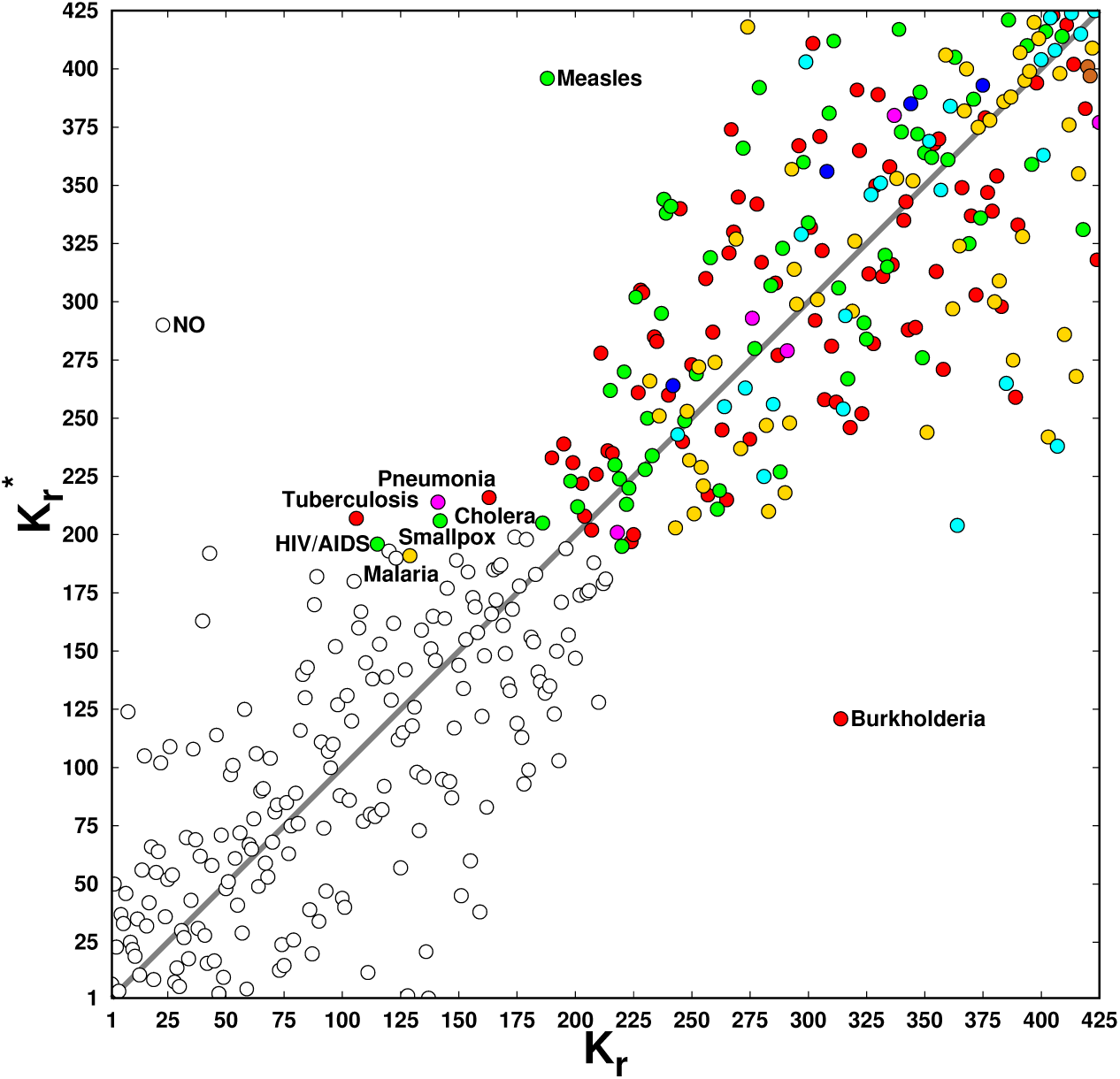
Distribution of the articles devoted to infectious diseases (colored circles) and to countries (white circles) in the PageRank ***K**_r_* – CheiRank 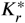 plane. The color code for infectious diseases is the same as in Fig. 1.

### Reduced Google matrix

To study further the selected subset of 425 nodes we use the reduced Google matrix approach and compute numerically *G*_R_ and its three components *G*_pr_, *G*_rr_, *G*_qr_. It is convenient to characterize each component by its weight defined as the sum of all elements divided by the matrix size *n_r_*. By definition we have the weight *W_R_* = 1 for *G*_R_ and we obtain weights *W_pr_* = 0.91021, *W_rr_* = 0.04715, *W_qr_* = 0.04264 (with nondiagonal weight *W_qrnd_* = 0.02667) respectively for *G*_pr_, *G*_rr_, *G*_qr_ (*G*_qrnd_). The weight of *G*_pr_ is signi1cantly larger than others but this matrix is close to the matrix composed from equal columns where the column is the PageRank vector (see also discussions in ***Frahm et al. (2016)***; ***El Zant et al. (2018a)*; *Lages et al. (2018)***). Due to this reason the components *G*_rr_ and *G*_qr_ provide an important information about interactions of nodes. Since the weights of these two components are approximately equal we see that the direct and indirect (hidden) links have a comparable contribution.

As an illustration we show in Fig. 5 a close up on African countries and viral diseases sectors of the full *n_r_* × *n_r_* reduced Google matrix *G*_R_ is shown (there are 55 African countries and 60 viral diseases shown in fig5). Detailed presentations of the *G*_R_ matrix components for the complete subset of *n_r_* = 425 countries and infectious diseases are given in ***Wiki4InfectiousDiseases (2018)***. In Fig. 5, *G*_R_ and its components are composed of diagonal blocks corresponding to country → country and disease → disease effective links, and off-diagonal blocks corresponding to disease → country (upper off-diagonal block) and country → disease (lower off-diagonal block) effective links.

**Figure 5.**
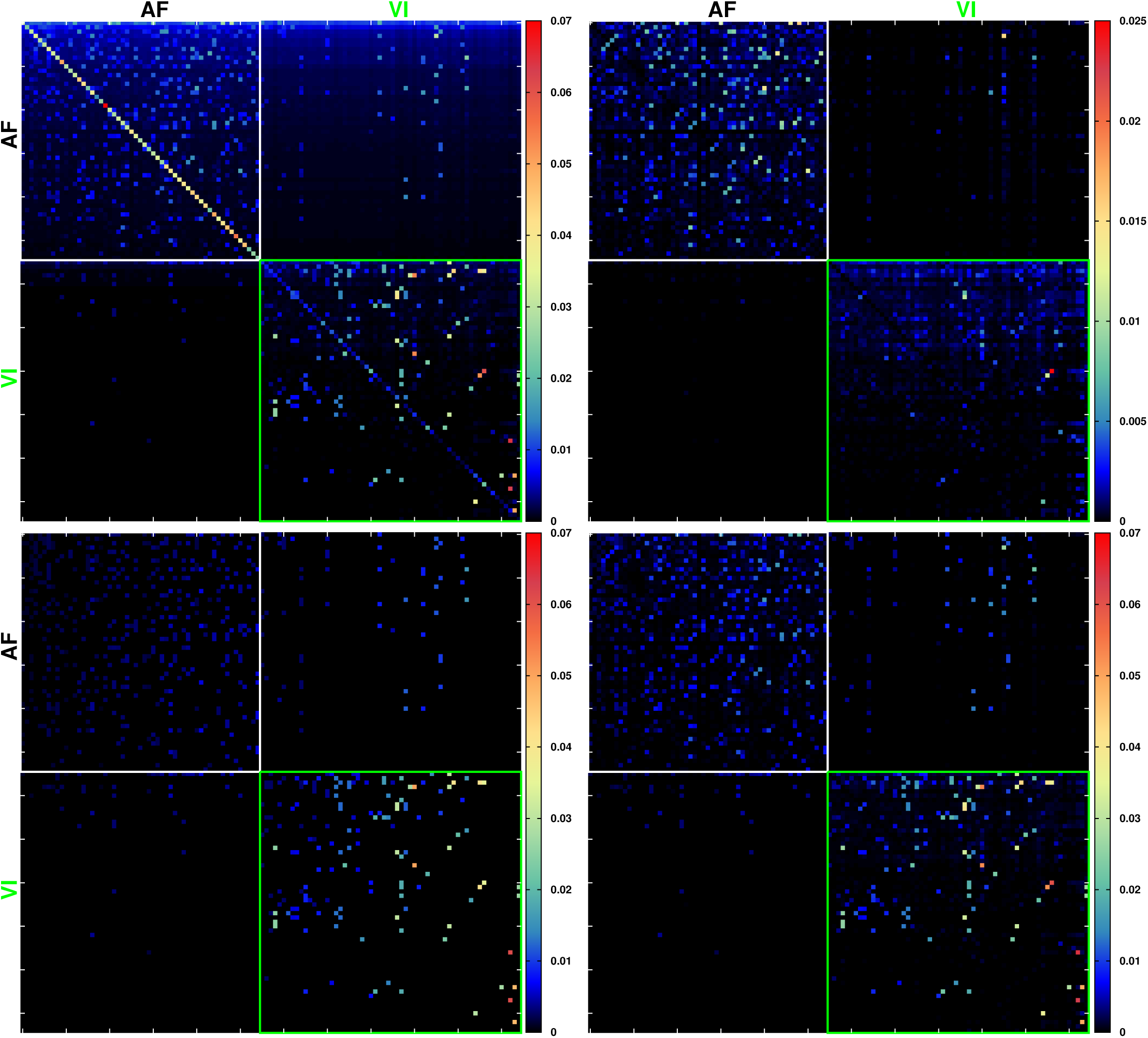
Reduced Google matrix *G*_R_ (top left panel) and three of its components, *G*_qrnd_ (top right panel), *G*_rr_ (bottom left panel), and *G*_rr_ + *G*_qrnd_, associated to the subnetwork of articles (bottom right) devoted to countries and infectious diseases in 2017 English Wikipedia. For the sake of clarity, here we show only matrix entries corresponding to the subset of articles devoted to African countries and viral diseases (see ***Wiki4InfectiousDiseases (2018)*** for the full subnetwork constituted by the *n_r_* = 425 articles devoted to all countries and all infectious diseases). Here lines and columns corresponding to African countries (viral diseases) are ordered as in Tab. 3 (Tab. 1). Horizontal and vertical tics on close-ups are placed every 10 entries.

### Friendship network of nodes

We use the matrix of direct and indirect transition *G*_rr_ + *G*_qr_ to determine the proximity relations between 230 nodes corresponding to diseases and to all *n_r_* = 425 nodes of diseases and countries. We call this the friendship networks being shown in Fig. 6. For each of 7 disease groups (see Tab. 2) we take a group leader as a disease with highest PageRank probability inside the group. Then on each step (level) we take 2 best friends define them as those nodes to which a leader has two highest transition matrix elements of *G*_rr_ + *G*_qr_. This gives us the second level of nodes below the 7 leaders. After that we generate the third level keeping again two better friends of the nodes of second level (those with highest transition probabilities). This algorithm is repeated until no new friends are found and the algorithm stops. In this way we obtain the network of 17 infectious diseases shown in the top panel of Fig. 6. The full arrows show the proximity links between disease nodes. The red arrows mark links with dominant contribution of *G*_qr_ indirect transitions while the black ones mark the links with dominance of *G*_rr_ direct transitions. Full arrows are for transitions from group leaders to nodes of second level, etc (see Fig. 6 caption for details). The obtained network is drawn with the Cytoscape software (***Shannon et al., 2003***).

**Figure 6.**
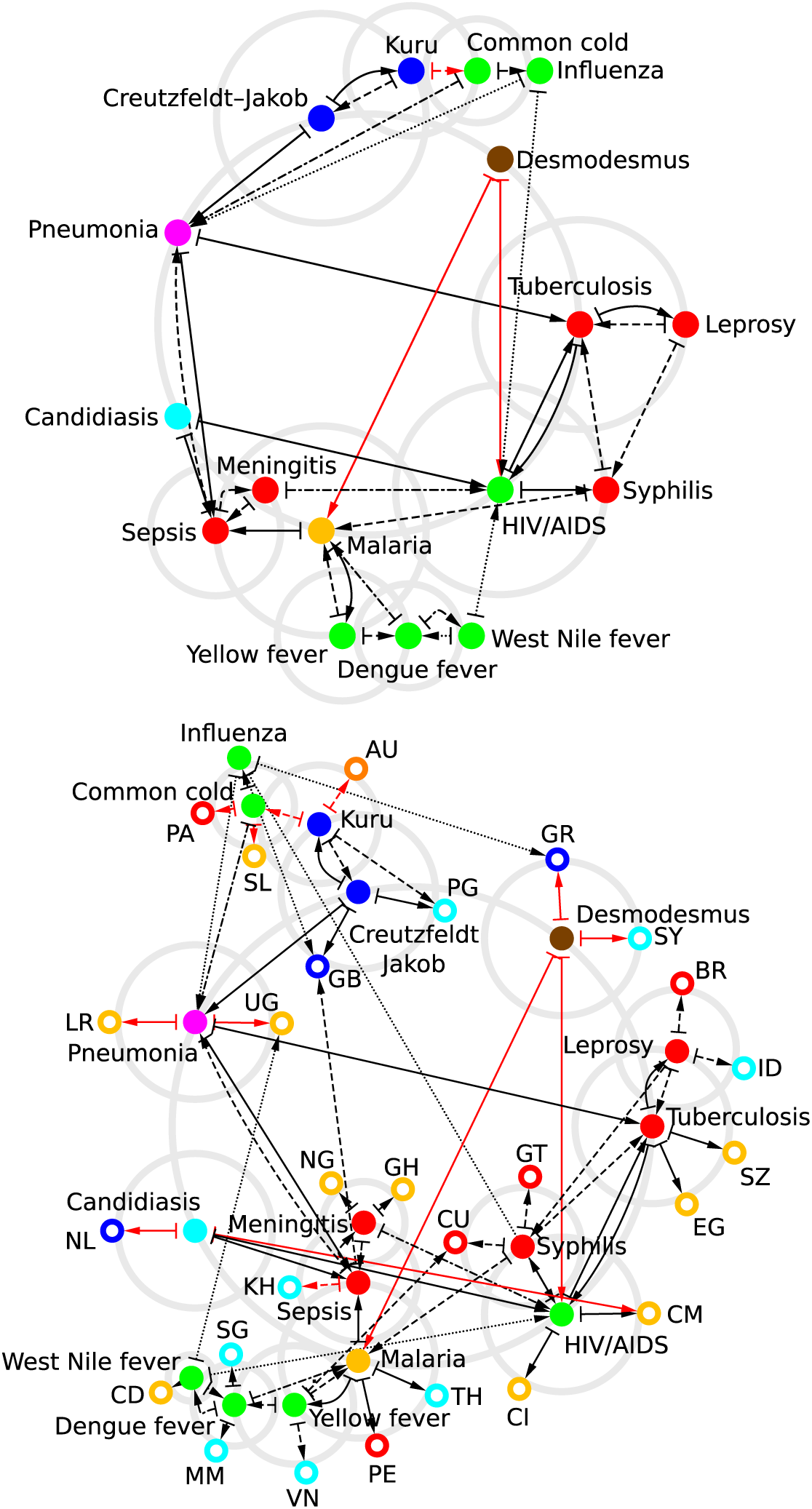
Infectious diseases friendship network. Top panel: we consider the set of top PageRank infectious disease for each of the seven type of infectious diseases, i.e., tuberculosis for bacterial type, HIV/AIDS for virus type, malaria for parasitic type, candidiasis for fungal type, pneumonia for multiple type, Creutzfeldt–Jakob for prionic type, and Desmodesmus for other type (see Tab. 1). From each one of these top PageRank diseases (placed along the main grey circle) we determine the two best linked diseases in *G*_rr_ + *G*_qr_, e.g., from tuberculosis the two best linked diseases are leprosy and HIV/AIDS. If not already present in the reduced network, we add these best linked diseases along secondary circles centered on the previous diseases. Then from each one of the new added diseases we determine the two best linked diseases, and so on. At the fourth iterations of this process no new diseases can be added. The arrows represent the links between diseases (1st iteration: plain line; 2nd iteration: dashed line; 3rd iteration: dashed–dotted line; 4th iteration: dotted line). Black arrows correspond to links existing in the adjacency matrix, red arrows are purely hidden links absent from adjacency matrix but present in *G*_qr_ component of the reduced Google matrix *G*_R_. The color code for infectious diseases is the same as in Fig. 1. Bottom panel: same reduced network as in the top panel but at each iteration also the best linked countries are determined. At each iteration no new links are determined from the newly added countries. Countries are represented by ring shape nodes with red color for countries from Americas, gold for African countries, cyan for Asian countries, blue for European countries, and orange for Oceanian countries. Network drawn with Cytoscape (***Shannon et al., 2003***).

In Fig. 6 top panel, four of the leader diseases are well connected; nodes corresponding to Tuberculosis (bacterial disease), HIV/AIDS (viral disease), Malaria (parasitic disease), Pneumonia (multiple origin disease) have 6 or more degrees. Nodes of Creutzfeldt-Jakob disease (prion disease) and of Desmodesmus (other origin disease) are more isolated. Focusing on first level friendship links (solid arrows), we retrieve several well known interactions between infectious diseases such as:

- the Tuberculosis–HIV/AIDS syndemic (see e.g. ***Pawlowski et al., 2012*; *Kwan and Ernst, 2011***) which is here represented by a closed loop between the two diseases,
- the interaction between HIV/AIDS and Syphilis (see e.g. ***Karp et al., 2009***) which is a typical example of syndemic between AIDS and sexually transmitted diseases,
- the Candidiasis interaction with HIV/AIDS, since the former is a very common opportunistic fungal infection for patients with HIV/AIDS (see e.g. ***Armstrong-James et al., 2014***), and the Candidiasis interaction with Sepsis1 since e.g. invasive Candidiasis which in some rare cases can lead to fulminant sepsis with an associated mortality exceeding 70% (see e.g. ***Pappas et al., 2018***),
- the Pneumonia to Sepsis interaction or the Malaria to Sepsis interaction, the first interaction reflects the fact that Sepsis is one of the possible complications of Pneumonia, the second interaction reflects that symptoms of Malaria resemble to those of Sepsis ***Auma et al. (2013)***,
- the closed loop interaction between Tuberculosis and Leprosy reflecting that these two diseases are caused by two different species of mycobacteria (see e.g. ***Wikipedia, 2018e***),
- the relation between Pneumonia and Tuberculosis, two severe pulmonary diseases (see e.g. ***WHO, 2018***),
- the Creutzfeldt–Jakob disease pointing to Pneumonia since patient infected by this prion disease develop a fatal Pneumonia due to impaired coughing reflexes (see e.g. ***Al Balushi et al., 2016***),
- the closed loop interaction between Kuru and Creutzfeldt–Jakob diseases since these two diseases are representatives of transmissible spongiform encephalopathies (see e.g. ***Sikorska and Liberski, 2012***). Taking into account also the other 2nd to 4th friendship levels, peculiar features appear such as:
- the cluster of bacterial diseases Tuberculosis–Leprosy–Syphilis; since there were a confusion between Leprosy and Syphilis in diagnosis before XXth century (see e.g. ***Leprosy and Syphilis, 1890***; ***Syphilis and Leprosy, 1899***), and false positives can occur with Tuberculosis for patients with Syphilis (see e.g. ***Shane, 2006***),
- the mosquito diseases cluster grouping Malaria, Yellow fever, Dengue fever and West Nile fever,
- the Meningitis–Sepsis closed loop since the Sepsis is usually developed at early stage by patient with Meningitis (see e.g. ***CDC, 2018***).

Red arrows in Fig. 6 indicate pure indirect links between infectious diseases: Desmodesmus and Malaria are both waterborne diseases (see e.g. ***Wikipedia, 2018f***), Desmodesmus and HIV/AIDS are related by a Wikipedia page devoted to immunocompetence (see e.g. ***Wikipedia, 2018b***), and Kuru in Papua New Guinea Foré language possibly means to shiver from cold (see e.g. ***Liberski and Ironside, 2015***).

From the above analysis we observe that the wiring between infectious diseases is meaningful guaranteeing that information encoded in the reduced Google matrix *G*_R_, and more precisely in its *G*_rr_ + *G*_qr_ component, is reliable, and can be used to infer possible relations between infectious diseases and any other subjects contained in Wikipedia such as e.g. countries, drugs, proteins, etc.

In the bottom panel of Fig. 6 we analyze the proximity between diseases of top panel with the world countries. Thus we add the two better “friend” countries being those to which a given disease has most strong matrix elements in *G*_rr_ + *G*_qr_ (there is no next iterations for country nodes). The friend countries (or proximity countries) are Egypt and Swaziland for Tuberculosis; Cameroon and Cote d’Ivoire for HIV/AIVS; Peru and Thailand for Malaria; Liberia and Uganda for Pneumonia; United Kingdom (UK)^2^ and Papua New Guinea for Creutzfeldt-Jakob disease and others. These strong links from an infectious disease to a given country well correspond to known events involving a disease and a country, like e.g. UK and Creutzfeldt-Jakob disease. We will see this in a more direct way using the sensitivity analysis presented in the next subsection.

### World country sensitivity to infectious diseases

We also preform analysis of the sensitivity *D*(*d* → *c, c*’) of country node *c*’ to the variation of the link *d* → *c*, where *d* denotes a disease node and *c* a country node. The diagonal sensitivity *D*(*d* → *c, c*) of world countries to Tuberculosis and HIV/AIDS are shown in Fig. 7. The most sensitive countries to Tuberculosis are Swaziland (SZ), Egypt (EG) and New Zealand (NZ). Indeed, in 2007 SZ had the highest estimated incidence rate of Tuberculosis as it is described in the corresponding Wikipedia article. Egypt also appears in this article since tubercular decay has been found in the spine of Egyptian mummy kept in the British Museum. NZ is present in this article since this country had a relatively successful effort to eradicate bovine tuberculosis. Thus Tuberculosis has direct links to these three countries (in agreement with two close country friends shown in the network of Fig. 6) that results in their high sensitivity to this disease. Of course, the origins of this sensitivity are different for SZ, EG, NZ. Thus Wikipedia network integrates all historical events related to Tuberculosis including ancient Egyptian mummy and recent years of high incidence rate in SZ. It can be discussed how important are these rather different types of links between disease and counties. Of course, a simplified network view cannot take into account all richness of historical events and describe them by a few number of links. However this approach provides a reliable global view of the interactions and dependencies between a disease and world countries.

**Figure 7.**
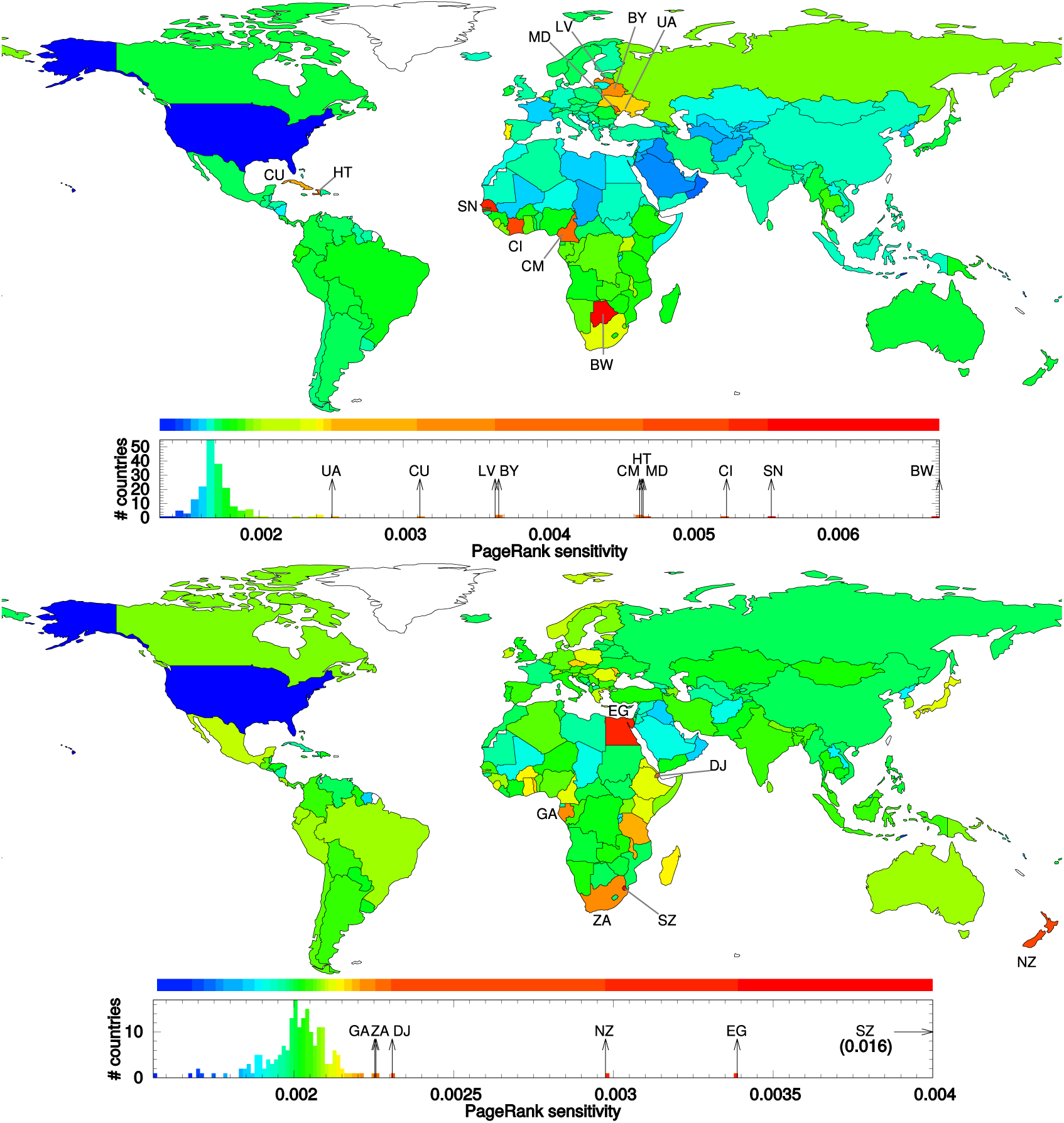
Country PageRank sensitivity to the variation of the reduced Google matrix HIV→country link (upper panel) and to the variation of the reduced Google matrix Tuberculosis→country link (bottom panel). The color categories are obtained using the Jenks natural breaks classi1cation method (***Wikipedia, 2018c***).

The sensitivity of countries to HIV/AIDS is shown in the top panel of Fig. 7. The most sensitive countries are Botswana (BW), Senegal (SN) and Cote d’Ivoire (CI). This happens since HIV/AIDS article directly points that estimated life expectancy in BW dropped from 65 to 35 years in 2006; SN and CI appears since the closest relative of HIV-2 exists in monkey living in coastal West Africa from SN to CI. The friendship network in Fig. 6 indeed marks the countries close to HIV/AIDS as CI and Cameroon (CM). The sensitivity map in Fig. 7 also shows that CM has high sensitivity to HIV/AIDS since HIV-1 appears to have originated in southern Cameroon.

The case of two diseases considered in Fig. 7 demonstrates that the REGOMAX approach is able to reliably determine the sensitivity of world countries to infectious diseases taking into account their relations on a scale of about 3 thousands of years.

A part of the sensitivity relations between diseases and countries can be visible from the friendship network as those in the bottom panel of Fig. 6. However, the REGOMAX approach can handle also indirect sensitivity *D*(*d* → *c, c*’) which is rather hard to be directly extracted from the friendship network. The examples of indirect sensitivity are shown in Fig. 8. Thus the variation of link from HIV/AIDS to Cameroon (CM) (Fig. 8 bottom panel) mainly affects Equatorial Guinea (GQ), Central African Republic (CF) and Chad (TD). The variation of link HIV/AIDS to USA (Fig. 8 top panel) produces the strongest sensitivity for Federal States of Micronesia (FM), Marshall Islands (MH) and Rwanda (RW). These countries are not present in the Wikipedia article HIV/AIDS and the obtained sensitivity emerges from a complex network interconnections between HIV/AIDS, USA (or Cameroon) to these countries. Thus the REGOMAX analysis allows to recover all network complexity of direct and indirect interactions between nodes.

**Figure 8.**
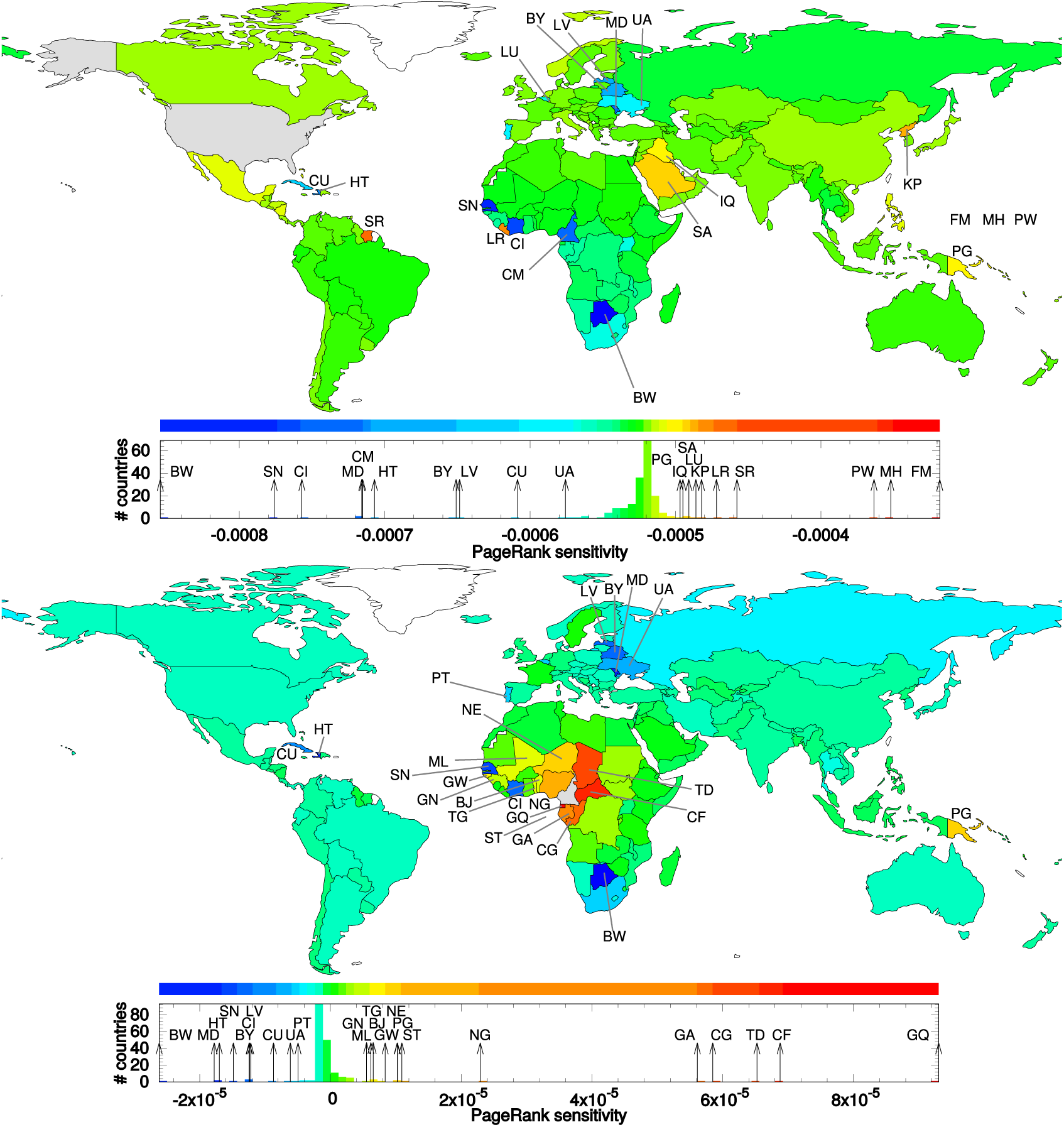
Country PageRank sensitivity to the variation of the reduced Google matrix HIV → USA link (top panel) and to the variation of the reduced Google matrix HIV → Cameroon link (bottom panel). The color categories are obtained using the Jenks natural breaks classi1cation method (***Wikipedia, 2018c***).

### Comparison of REGOMAX and WHO results

It is important to compare the results of REGOMAX analysis with those of World Health Organization (WHO) or other sources on number of infected people. With this aim we extract from WHO reports (***WHO, 2008*, *2018***) the number of Tuberculosis incidences per 100000 population of a given country, and the number of new HIV/AIDS infections per 1000 uninfected population. The ranking of countries by the number of incidences is presented in Tab. 4 for Tuberculosis and the ranking of countries by new infections is presented in Tab. 5 for HIV/AIDS. We also analyze WHO 2004 data (***WHO, 2008***) for the number of deaths caused by each disease in 2004 complemented by the data reported in ***Wikipedia (2018a)***. The resulting ranking of diseases is given in Tab. 6.

We compare these official WHO ranking results with those obtained from REGOMAX analysis. Thus we determine the ranking of countries by their sensitivity to Tuberculosis and to HIV/AIDS for top 100 countries (these ranking lists are given in ***Wiki4InfectiousDiseases (2018)***). The overlap *η*(*j*) of these REGOMAX rankings with those of WHO from Tab. 4 and Tab. 5 are shown in Fig. 9. We obtain the overlap of 50% for Tuberculosis and 79% for HIV/AIDS for the top 100 countries. These numbers are comparable with overlaps obtained for top 100 historical figures found from Wikipedia and historical analysis (see ***Eom et al. (2015)***) and for top 100 world universities determined by Wikipedia and Shanghai ranking (see ***Lages et al. (2016)*; *Coquidé et al. (2018)***).

**Figure 9.**
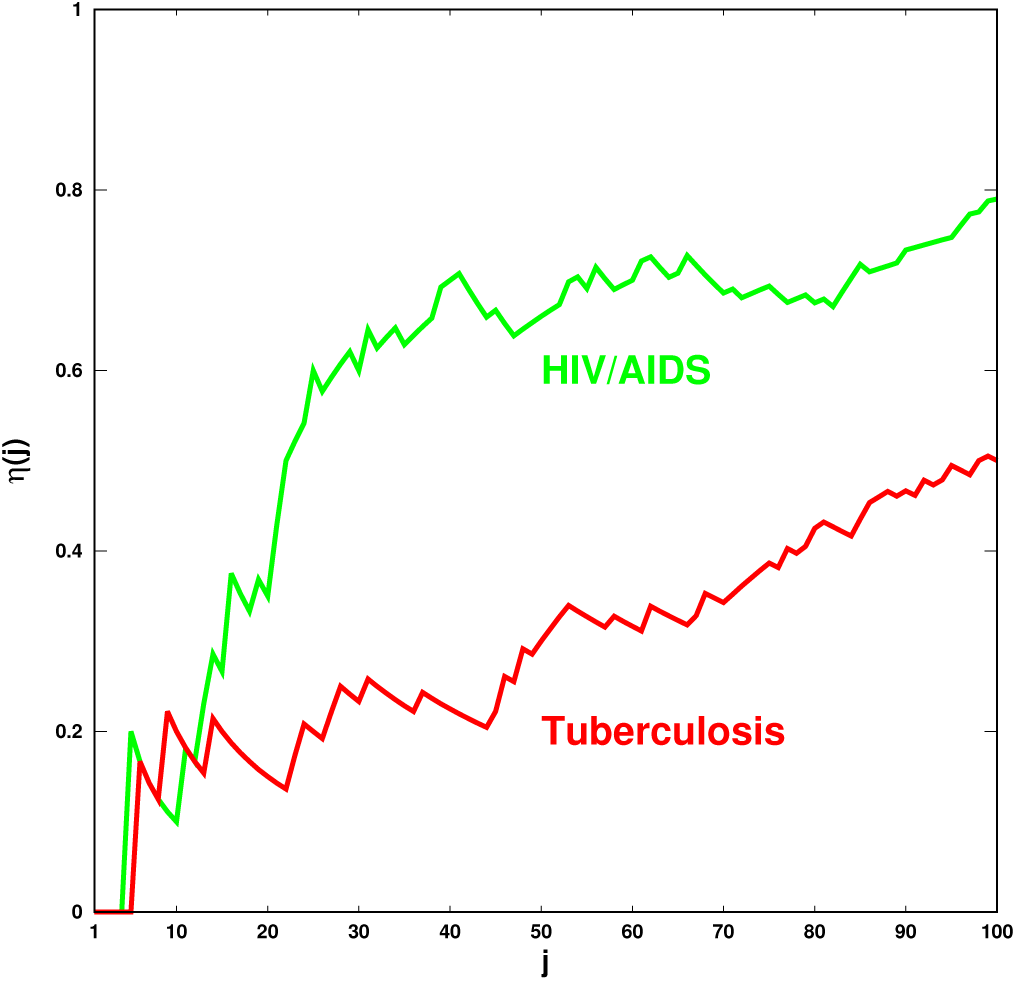
Overlap *η*(*j*) = *j_c_* /*j* between the ranking of countries obtained from the REGOMAX PageRank sensitivity computation and the ranking of countries obtained from WHO data (Tab. 4 for Tuberculosis and Tab. 5 for HIV/AIDS). Here, *j_c_* is the number of common countries in the top *j* of the two rankings. Green curve is for the HIV/AIDS disease, and red curve is for the Tuberculosis disease.

Another comparison is presented in Fig. 10. Here we take the infectious diseases ordered by their PageRank index and compare them with the ranking list of diseases ordered by the number of deaths caused by them in 2004 (see Tab. 6). The obtained overlap is shown in Fig. 10 with an overall of 100% for top 4 deadliest diseases (which from Tab. 6 are 1 Pneumonia, 2 HIV/AIDS, 3 Tuberculosis, 4 Malaria) and 54% for the whole list of 31 considered diseases. In addition the REGOMAX analysis allows to determine the world map of countries with the highest sensitivity to the top list of 7 diseases. Such maps are shown in Fig. 11. We see the world dominance of Tuberculosis (top panel) while after it (or in its absence) we 1nd the world dominance of HIV/AIDS (bottom panel). The geographical influence of Malaria and Diarrheal diseases are also well visible.

**Figure 10.**
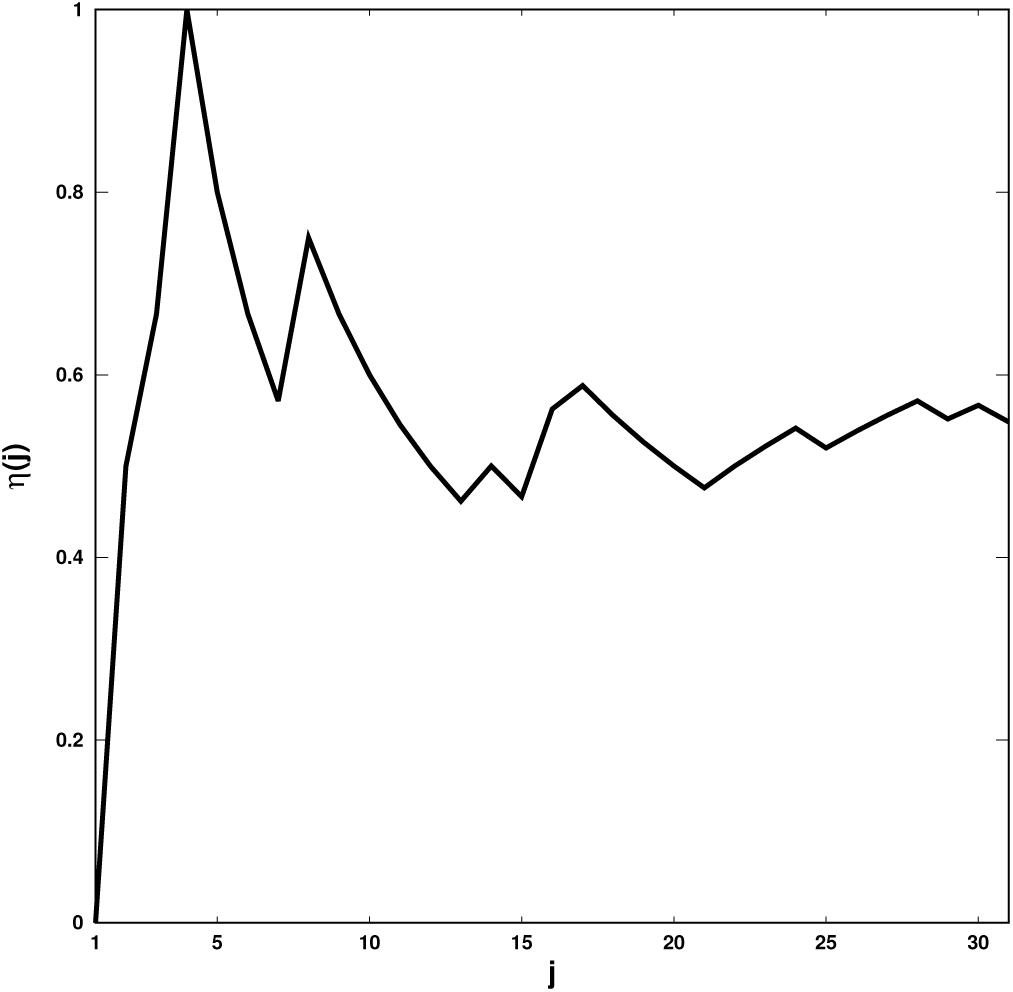
Overlap *η*(*j*) = *j_c_* /*j* between the PageRank of the diseases and the diseases ranking based on the number of deaths during the year 2004 (Tab. 6). Here, *j_c_* is the number of common countries in the top *j* of the two rankings. The numbers of deaths caused by each disease during the year 2004 are extracted from ***WHO (2008)***. For 8 of these diseases, the number of deaths have been estimated using English Wikipedia edition (***Wikipedia, 2018a***).

**Figure 11.**
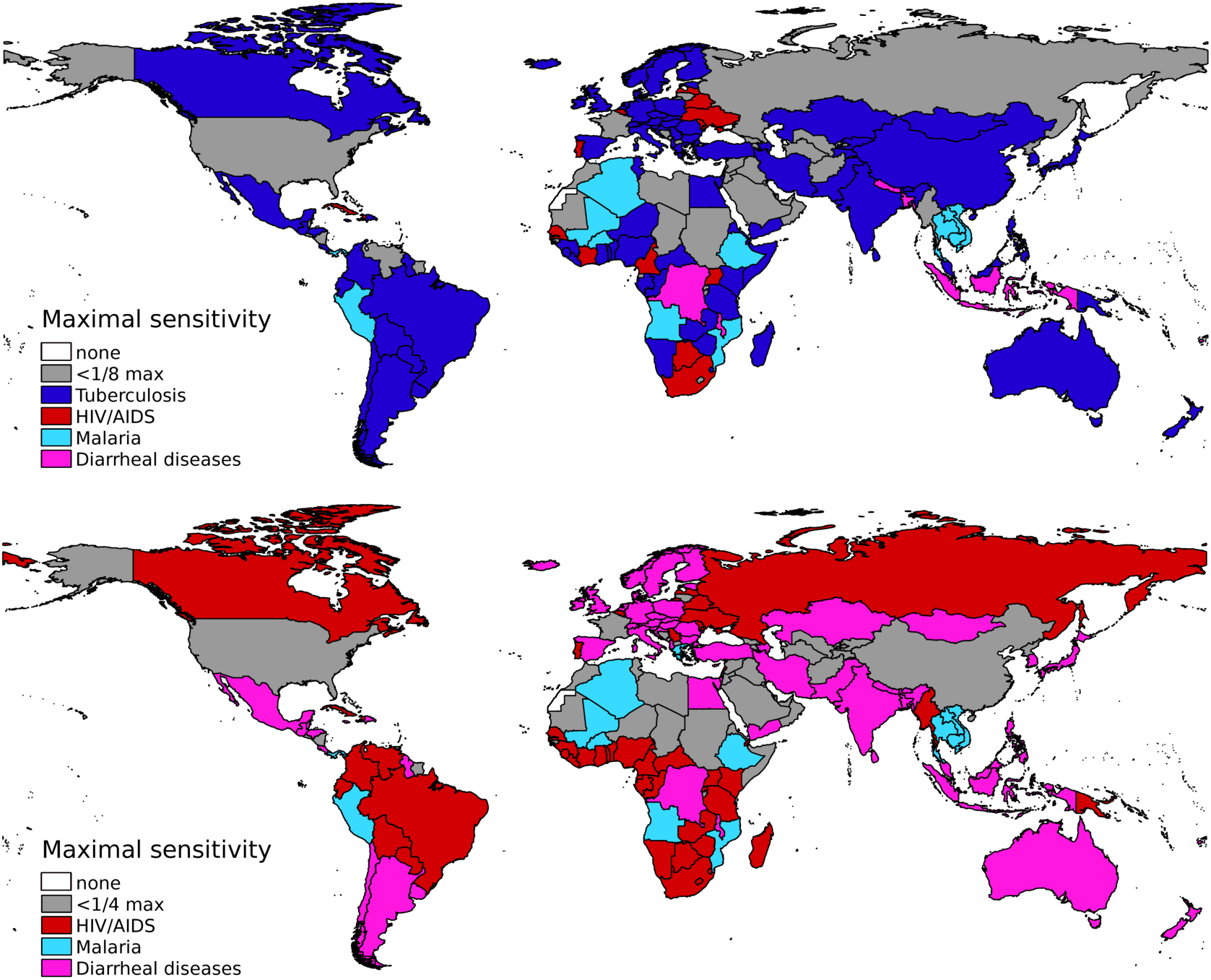
Maps of the maximal sensitivity of diseases. Each country is colored according to the disease giving the maximal sensitivity: Tuberculosis (blue), HIV/AIDS (red), Malaria (cyan), Diarrheal diseases (magenta), Dengue, Meningitis, Hepatitis. Top panel shows the maximal sensitivity for the 7 diseases, and bottom panel for 6 of these diseases (Tuberculosis is not taken into account). Gray color indicates countries for which the maximal sensitivity to these diseases is less than 1/8 (1/4) of the greatest maximal sensitivity for any country. Note that Dengue, Meningitis and Hepatitis never gives the maximal sensitivity for any country.

Thus the performed comparison with WHO data shows that the Google matrix analysis of Wikipedia network provides us a reliable information about world importance and influence of infectious diseases.

## Discussion

In this work we presented the reduced Google matrix (or REGOMAX) analysis of world influence of infectious diseases from the English Wikipedia network of 2017. This method allows to take into account all direct and indirect links between the selected nodes of countries and diseases. The importance of diseases is determined by their PageRank probabilities. The REGOMAX analysis allows to establish the network of proximity (friendship) relations between the diseases and countries. The sensitivity of world countries to a specific disease is determined as well as the influence of link variation between a disease and a country on other countries. The comparison with the WHO data confirms the reliability of REGOMAX results applied to Wikipedia network.

**Appendix 1 Table 1.**
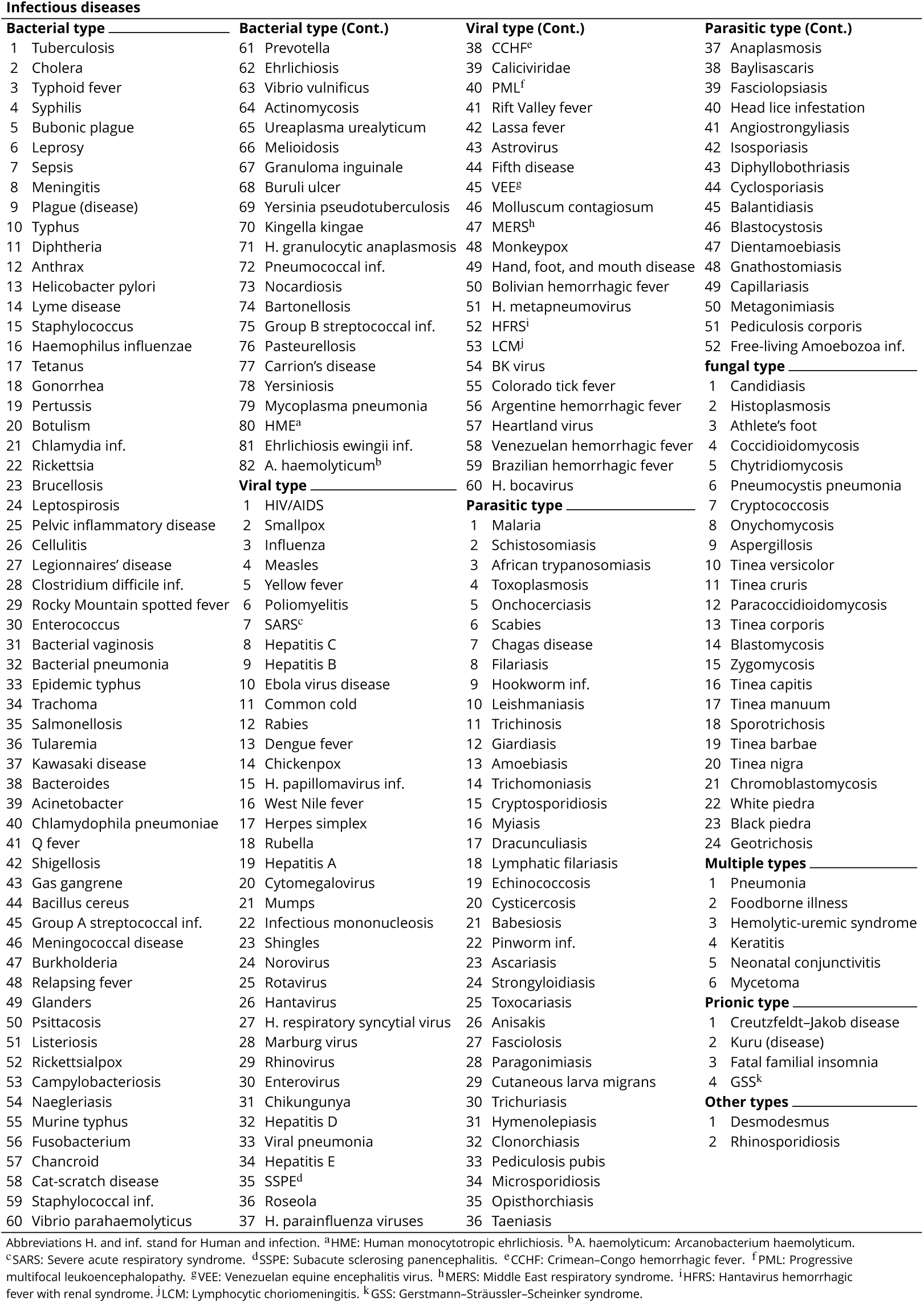
List of infectious diseases ordered by type (82 of bacterial type, 60 of viral type, 52 of parasitic type, 24 of fungal type, 6 of multiple types, and 2 of other types) then by PageRank of the corresponding article in English edition of Wikipedia (***Frahm and Shepelyansky, 2017***). The *n_d_* = 230 elements of this list have been extracted from the *list of infectious diseases* article in 2017 English Wikipedia (***Wikipedia, 2018d*; *Wiki4InfectiousDiseases, 2018***).

**Appendix 1 Table 2.**
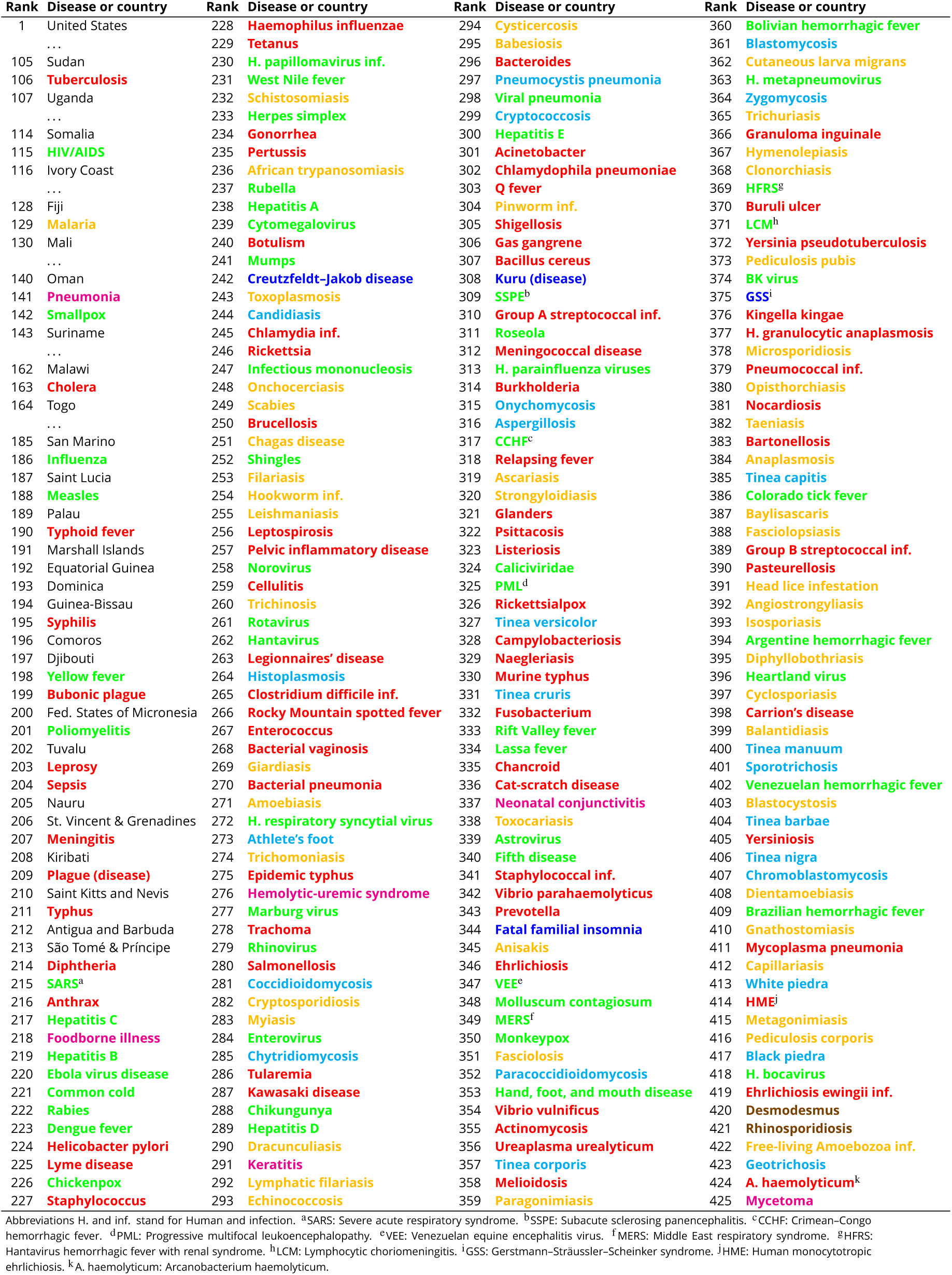
Ranking of infectious diseases and countries in English Wikipedia 2017 according to PageRank algorithm. The color code distinguishes type of infectious diseases: bacterial, viral, parasitic, fungal, prionic, multiple origin, and other origin.

**Appendix 1 Table 3.** List of countries ordered by PageRank of the corresponding article in English Wikipedia 2017. Here the countries correspond to the *n_c_* = 195 sovereign states listed in (***Wikipedia, 2017***). The country code (CC) is given for each country.

| Rank | Country | CC | Rank | Country | CC | Rank | Country | CC |
| --- | --- | --- | --- | --- | --- | --- | --- | --- |
| 1 | United States | US | 66 | Morocco | MA | 131 | Kyrgyzstan | KG |
| 2 | France | FR | 67 | Cuba | CU | 132 | Guyana | GY |
| 3 | Germany | DE | 68 | Algeria | DZ | 133 | Trinidad and Tobago | TT |
| 4 | United Kingdom | GB | 69 | Bosnia and Herzegovina | BA | 134 | Mauritius | MU |
| 5 | Iran | IR | 70 | Ecuador | EC | 135 | Tajikistan | TJ |
| 6 | India | IN | 71 | Saudi Arabia | SA | 136 | Monaco | MC |
| 7 | Canada | CA | 72 | Lithuania | LT | 137 | Oman | OM |
| 8 | Australia | AU | 73 | Iceland | IS | 138 | Suriname | SR |
| 9 | China | CN | 74 | Bolivia | BO | 139 | Liberia | LR |
| 10 | Italy | IT | 75 | Tanzania | TZ | 140 | Solomon Islands | SB |
| 11 | Japan | JP | 76 | Ethiopia | ET | 141 | Sierra Leone | SL |
| 12 | Russia | RU | 77 | Democratic Republic of the Congo | CD | 142 | Bhutan | BT |
| 13 | Brazil | BR | 78 | Madagascar | MG | 143 | The Bahamas | BS |
| 14 | Spain | ES | 79 | Armenia | AM | 144 | Vatican City | VA |
| 15 | Netherlands | NL | 80 | Lebanon | LB | 145 | Bahrain | BH |
| 16 | Poland | PL | 81 | Cyprus | CY | 146 | Barbados | BB |
| 17 | Sweden | SE | 82 | Kazakhstan | KZ | 147 | Botswana | BW |
| 18 | Mexico | MX | 83 | Georgia (country) | GE | 148 | Rwanda | RW |
| 19 | Turkey | TR | 84 | Latvia | LV | 149 | Turkmenistan | TM |
| 20 | Romania | RO | 85 | Panama | PA | 150 | Benin | BJ |
| 21 | New Zealand | NZ | 86 | Belarus | BY | 151 | Niger | NE |
| 22 | South Africa | ZA | 87 | Albania | AL | 152 | Gabon | GA |
| 23 | Norway | NO | 88 | Papua New Guinea | PG | 153 | Brunei | BN |
| 24 | Switzerland | CH | 89 | Luxembourg | LU | 154 | Belize | BZ |
| 25 | Philippines | PH | 90 | Ghana | GH | 155 | Guinea | GN |
| 26 | Austria | AT | 91 | United Arab Emirates | AE | 156 | Chad | TD |
| 27 | Belgium | BE | 92 | Uruguay | UY | 157 | Malawi | MW |
| 28 | Pakistan | PK | 93 | North Korea | KP | 158 | Togo | TG |
| 29 | Argentina | AR | 94 | Yemen | YE | 159 | Liechtenstein | LI |
| 30 | Indonesia | ID | 95 | Costa Rica | CR | 160 | Samoa | WS |
| 31 | Greece | GR | 96 | Malta | MT | 161 | Burundi | BI |
| 32 | Denmark | DK | 97 | Tunisia | TN | 162 | South Sudan | SS |
| 33 | South Korea | KR | 98 | Jamaica | JM | 163 | Republic of the Congo | CG |
| 34 | Israel | IL | 99 | Zimbabwe | ZW | 164 | East Timor | TL |
| 35 | Hungary | HU | 100 | Cambodia | KH | 165 | Cape Verde | CV |
| 36 | Finland | FI | 101 | Cameroon | CM | 166 | Eritrea | ER |
| 37 | Egypt | EG | 102 | Mongolia | MN | 167 | Mauritania | MR |
| 38 | Portugal | PT | 103 | Burkina Faso | BF | 168 | Central African Republic | CF |
| 39 | Ukraine | UA | 104 | Jordan | JO | 169 | Maldives | MV |
| 40 | Sri Lanka | LK | 105 | Sudan | SD | 170 | Tonga | TO |
| 41 | Czech Republic | CZ | 106 | Uganda | UG | 171 | Andorra | AD |
| 42 | Malaysia | MY | 107 | Republic of Macedonia | MK | 172 | Vanuatu | VU |
| 43 | Peru | PE | 108 | Guatemala | GT | 173 | State of Palestine | PS |
| 44 | Thailand | TH | 109 | Libya | LY | 174 | Lesotho | LS |
| 45 | Colombia | CO | 110 | Dominican Republic | DO | 175 | The Gambia | GM |
| 46 | Bulgaria | BG | 111 | Haiti | HT | 176 | Swaziland | SZ |
| 47 | Chile | CL | 112 | Moldova | MD | 177 | Seychelles | SC |
| 48 | Republic of Ireland | IE | 113 | Somalia | SO | 178 | Grenada | GD |
| 49 | Singapore | SG | 114 | Ivory Coast | CI | 179 | San Marino | SM |
| 50 | Serbia | RS | 115 | Namibia | NA | 180 | Saint Lucia | LC |
| 51 | Azerbaijan | AZ | 116 | Paraguay | PY | 181 | Palau | PW |
| 52 | Vietnam | VN | 117 | Angola | AO | 182 | Marshall Islands | MH |
| 53 | Nepal | NP | 118 | Uzbekistan | UZ | 183 | Equatorial Guinea | GQ |
| 54 | Estonia | EE | 119 | Montenegro | ME | 184 | Dominica | DM |
| 55 | Croatia | HR | 120 | Kuwait | KW | 185 | Guinea-Bissau | GW |
| 56 | Nigeria | NG | 121 | Laos | LA | 186 | Comoros | KM |
| 57 | Afghanistan | AF | 122 | Mozambique | MZ | 187 | Djibouti | DJ |
| 58 | Iraq | IQ | 123 | Nicaragua | NI | 188 | Federated States of Micronesia | FM |
| 59 | Bangladesh | BD | 124 | Qatar | QA | 189 | Tuvalu | TV |
| 60 | Syria | SY | 125 | Senegal | SN | 190 | Nauru | NR |
| 61 | Myanmar | MM | 126 | Fiji | FJ | 191 | Saint Vincent and the Grenadines | VC |
| 62 | Kenya | KE | 127 | Mali | ML | 192 | Kiribati | KI |
| 63 | Slovakia | SK | 128 | Honduras | HN | 193 | Saint Kitts and Nevis | KN |
| 64 | Venezuela | VE | 129 | Zambia | ZM | 194 | Antigua and Barbuda | AG |
| 65 | Slovenia | SI | 130 | El Salvador | SV | 195 | São Tomé and Príncipe | ST |

**Appendix 1 Table 4.** List of countries ordered by Tuberculosis incidences in 2016 (per 100000 population per year; taken from ***WHO (2018)***)

| Rank | Country | Incidence | Rank | Country | Incidence | Rank | Country | Incidence |
| --- | --- | --- | --- | --- | --- | --- | --- | --- |
| 1 | South Africa | 781.0 | 65 | Mauritania | 102.0 | 129 | Portugal | 20.0 |
| 2 | Lesotho | 724.0 | 66 | Moldova | 101.0 | 130 | Serbia | 19.0 |
| 3 | Kiribati | 566.0 | 67 | São Tomé and Príncipe | 99.0 | 131 | Turkey | 18.0 |
| 4 | Philippines | 554.0 | 68 | Niger | 93.0 | 132 | Trinidad and Tobago | 18.0 |
| 5 | Mozambique | 551.0 | 69 | Guyana | 93.0 | 133 | Poland | 18.0 |
| 6 | North Korea | 513.0 | 70 | Malaysia | 92.0 | 134 | Montenegro | 16.0 |
| 7 | East Timor | 498.0 | 71 | Georgia (country) | 92.0 | 135 | Japan | 16.0 |
| 8 | Gabon | 485.0 | 72 | Ukraine | 87.0 | 136 | Estonia | 16.0 |
| 9 | Namibia | 446.0 | 73 | Tajikistan | 85.0 | 137 | Chile | 16.0 |
| 10 | Papua New Guinea | 432.0 | 74 | Solomon Islands | 84.0 | 138 | Albania | 16.0 |
| 11 | Marshall Islands | 422.0 | 75 | Sudan | 82.0 | 139 | Seychelles | 15.0 |
| 12 | Central African Republic | 407.0 | 76 | South Korea | 77.0 | 140 | Iran | 14.0 |
| 13 | Swaziland | 398.0 | 77 | Uzbekistan | 76.0 | 141 | Egypt | 14.0 |
| 14 | Indonesia | 391.0 | 78 | Romania | 74.0 | 142 | Malta | 13.0 |
| 15 | Republic of the Congo | 378.0 | 79 | Eritrea | 74.0 | 143 | Lebanon | 12.0 |
| 16 | Zambia | 376.0 | 80 | Algeria | 70.0 | 144 | Croatia | 12.0 |
| 17 | Guinea-Bissau | 374.0 | 81 | Kazakhstan | 67.0 | 145 | Bahrain | 12.0 |
| 18 | Angola | 370.0 | 82 | Russia | 66.0 | 146 | Spain | 10.0 |
| 19 | Myanmar | 361.0 | 83 | Brunei | 66.0 | 147 | Saudi Arabia | 10.0 |
| 20 | Kenya | 348.0 | 84 | Azerbaijan | 66.0 | 148 | Belgium | 10.0 |
| 21 | Cambodia | 345.0 | 85 | Sri Lanka | 65.0 | 149 | United Kingdom | 9.9 |
| 22 | Djibouti | 335.0 | 86 | China | 64.0 | 150 | Costa Rica | 9.5 |
| 23 | Botswana | 326.0 | 87 | Turkmenistan | 60.0 | 151 | Oman | 9.0 |
| 24 | Democratic Republic of the Congo | 323.0 | 88 | El Salvador | 60.0 | 152 | Hungary | 8.8 |
| 25 | Liberia | 308.0 | 89 | Dominican Republic | 60.0 | 153 | Tonga | 8.6 |
| 26 | Sierra Leone | 304.0 | 90 | Fiji | 59.0 | 154 | Sweden | 8.2 |
| 27 | Tanzania | 287.0 | 91 | Benin | 59.0 | 155 | Austria | 8.2 |
| 28 | Somalia | 270.0 | 92 | Vanuatu | 56.0 | 156 | Germany | 8.1 |
| 29 | Pakistan | 268.0 | 93 | Mali | 56.0 | 157 | Switzerland | 7.8 |
| 30 | Madagascar | 237.0 | 94 | Panama | 55.0 | 158 | Dominica | 7.8 |
| 31 | Bangladesh | 221.0 | 95 | Lithuania | 53.0 | 159 | Samoa | 7.7 |
| 32 | Nigeria | 219.0 | 96 | Belarus | 52.0 | 160 | France | 7.7 |
| 33 | India | 211.0 | 97 | Singapore | 51.0 | 161 | New Zealand | 7.3 |
| 34 | Zimbabwe | 208.0 | 98 | Burkina Faso | 51.0 | 162 | Republic of Ireland | 7.1 |
| 35 | Tuvalu | 207.0 | 99 | Rwanda | 50.0 | 163 | Cuba | 6.9 |
| 36 | Cameroon | 203.0 | 100 | Ecuador | 50.0 | 164 | Slovenia | 6.5 |
| 37 | Uganda | 201.0 | 101 | Maldives | 49.0 | 165 | Grenada | 6.4 |
| 38 | Afghanistan | 189.0 | 102 | Yemen | 48.0 | 166 | Saint Vincent and the Grenadines | 6.3 |
| 39 | Haiti | 188.0 | 103 | Nicaragua | 48.0 | 167 | Norway | 6.1 |
| 40 | Mongolia | 183.0 | 104 | Togo | 46.0 | 168 | Italy | 6.1 |
| 41 | Equatorial Guinea | 181.0 | 105 | Armenia | 44.0 | 169 | Denmark | 6.1 |
| 42 | Bhutan | 178.0 | 106 | Iraq | 43.0 | 170 | Australia | 6.1 |
| 43 | Federated States of Micronesia | 177.0 | 107 | Paraguay | 42.0 | 171 | Andorra | 6.0 |
| 44 | Ethiopia | 177.0 | 108 | Brazil | 42.0 | 172 | Slovakia | 5.9 |
| 45 | Guinea | 176.0 | 109 | Libya | 40.0 | 173 | Netherlands | 5.9 |
| 46 | Laos | 175.0 | 110 | Honduras | 40.0 | 174 | Luxembourg | 5.8 |
| 47 | The Gambia | 174.0 | 111 | Tunisia | 38.0 | 175 | Jordan | 5.6 |
| 48 | Thailand | 172.0 | 112 | Belize | 38.0 | 176 | Cyprus | 5.6 |
| 49 | Malawi | 159.0 | 113 | Latvia | 37.0 | 177 | Canada | 5.2 |
| 50 | Ghana | 156.0 | 114 | Comoros | 35.0 | 178 | Czech Republic | 5.0 |
| 51 | Nepal | 154.0 | 115 | Venezuela | 32.0 | 179 | Finland | 4.7 |
| 52 | Ivory Coast | 153.0 | 116 | Colombia | 32.0 | 180 | Jamaica | 4.5 |
| 53 | Chad | 153.0 | 117 | Bosnia and Herzegovina | 32.0 | 181 | Greece | 4.4 |
| 54 | South Sudan | 146.0 | 118 | Uruguay | 29.0 | 182 | Israel | 3.5 |
| 55 | Kyrgyzstan | 145.0 | 119 | Bulgaria | 27.0 | 183 | Antigua and Barbuda | 3.4 |
| 56 | Senegal | 140.0 | 120 | Suriname | 26.0 | 184 | United States | 3.1 |
| 57 | Cape Verde | 137.0 | 121 | The Bahamas | 26.0 | 185 | Iceland | 2.1 |
| 58 | Vietnam | 133.0 | 122 | Kuwait | 24.0 | 186 | Saint Lucia | 1.9 |
| 59 | Palau | 123.0 | 123 | Guatemala | 24.0 | 187 | Barbados | 1.2 |
| 60 | Burundi | 118.0 | 124 | Argentina | 24.0 | 188 | United Arab Emirates | 0.79 |
| 61 | Peru | 117.0 | 125 | Qatar | 23.0 | 189 | San Marino | 0.0 |
| 62 | Bolivia | 114.0 | 126 | Mexico | 22.0 | 190 | Saint Kitts and Nevis | 0.0 |
| 63 | Nauru | 112.0 | 127 | Mauritius | 22.0 | 191 | Monaco | 0.0 |
| 64 | Morocco | 103.0 | 128 | Syria | 21.0 |  |  |  |

**Appendix 1 Table 5.** List of countries ordered by new HIV/AIDS infections in 2017 (per 1000 uninfected population; taken from ***WHO (2018)***)

| Rank | Country | New infections | Rank | Country | New infections | Rank | Country | New infections |
| --- | --- | --- | --- | --- | --- | --- | --- | --- |
| 1 | Lesotho | 12.68 | 44 | Ethiopia | 0.33 | 87 | Azerbaijan | 0.1 |
| 2 | Swaziland | 9.37 | 45 | Cape Verde | 0.31 | 88 | Spain | 0.09 |
| 3 | South Africa | 5.58 | 46 | Trinidad and Tobago | 0.29 | 89 | Peru | 0.09 |
| 4 | Botswana | 5.52 | 47 | Cuba | 0.29 | 90 | Niger | 0.09 |
| 5 | Namibia | 4.37 | 48 | Georgia (country) | 0.28 | 91 | Lithuania | 0.09 |
| 6 | Zambia | 4.08 | 49 | Chile | 0.28 | 92 | France | 0.09 |
| 7 | Mozambique | 3.63 | 50 | Dominican Republic | 0.24 | 93 | Armenia | 0.09 |
| 8 | Zimbabwe | 3.03 | 51 | Brazil | 0.24 | 94 | Senegal | 0.08 |
| 9 | Equatorial Guinea | 2.71 | 52 | Latvia | 0.23 | 95 | Albania | 0.08 |
| 10 | Malawi | 2.29 | 53 | Myanmar | 0.22 | 96 | Sweden | 0.06 |
| 11 | Central African Republic | 1.8 | 54 | Venezuela | 0.21 | 97 | Nicaragua | 0.06 |
| 12 | Republic of the Congo | 1.65 | 55 | Paraguay | 0.2 | 98 | Malta | 0.06 |
| 13 | Uganda | 1.5 | 56 | Burundi | 0.2 | 99 | Italy | 0.06 |
| 14 | Kenya | 1.46 | 57 | Malaysia | 0.19 | 100 | Republic of Ireland | 0.06 |
| 15 | Cameroon | 1.39 | 58 | Indonesia | 0.19 | 101 | Iran | 0.06 |
| 16 | South Sudan | 1.35 | 59 | Costa Rica | 0.19 | 102 | India | 0.06 |
| 17 | Nigeria | 1.23 | 60 | Burkina Faso | 0.19 | 103 | Australia | 0.05 |
| 18 | Tanzania | 1.19 | 61 | Madagascar | 0.18 | 104 | Yemen | 0.04 |
| 19 | Angola | 0.94 | 62 | Luxembourg | 0.18 | 105 | Romania | 0.04 |
| 20 | Gabon | 0.92 | 63 | Guatemala | 0.18 | 106 | Czech Republic | 0.04 |
| 21 | Sierra Leone | 0.86 | 64 | Somalia | 0.17 | 107 | Cambodia | 0.04 |
| 22 | Ivory Coast | 0.86 | 65 | Democratic Republic of the Congo | 0.17 | 108 | Bahrain | 0.04 |
| 23 | Ghana | 0.78 | 66 | Kazakhstan | 0.16 | 109 | Tunisia | 0.03 |
| 24 | Haiti | 0.77 | 67 | El Salvador | 0.16 | 110 | Sri Lanka | 0.03 |
| 25 | Guyana | 0.77 | 68 | Uruguay | 0.15 | 111 | Slovenia | 0.03 |
| 26 | Belize | 0.75 | 69 | Tajikistan | 0.15 | 112 | Serbia | 0.03 |
| 27 | Guinea-Bissau | 0.72 | 70 | Eritrea | 0.15 | 113 | Netherlands | 0.03 |
| 28 | Rwanda | 0.7 | 71 | Sudan | 0.13 | 114 | Nepal | 0.03 |
| 29 | Guinea | 0.67 | 72 | Kyrgyzstan | 0.13 | 115 | Morocco | 0.03 |
| 30 | Liberia | 0.66 | 73 | Argentina | 0.13 | 116 | Bulgaria | 0.03 |
| 31 | The Gambia | 0.65 | 74 | Vietnam | 0.12 | 117 | Afghanistan | 0.03 |
| 32 | Jamaica | 0.63 | 75 | Mauritania | 0.12 | 118 | Slovakia | 0.02 |
| 33 | Suriname | 0.62 | 76 | Fiji | 0.12 | 119 | Saudi Arabia | 0.02 |
| 34 | Togo | 0.59 | 77 | Ecuador | 0.12 | 120 | Qatar | 0.02 |
| 35 | Djibouti | 0.58 | 78 | Colombia | 0.12 | 121 | Lebanon | 0.02 |
| 36 | Barbados | 0.58 | 79 | Philippines | 0.11 | 122 | Kuwait | 0.02 |
| 37 | Ukraine | 0.38 | 80 | Montenegro | 0.11 | 123 | Egypt | 0.02 |
| 38 | Moldova | 0.38 | 81 | Honduras | 0.11 | 124 | Croatia | 0.02 |
| 39 | Papua New Guinea | 0.37 | 82 | Thailand | 0.1 | 125 | Algeria | 0.02 |
| 40 | Panama | 0.34 | 83 | Pakistan | 0.1 | 126 | Mongolia | 0.01 |
| 41 | Chad | 0.34 | 84 | Mexico | 0.1 | 127 | Jordan | <0.01 |
| 42 | Benin | 0.34 | 85 | Laos | 0.1 | 128 | Comoros | <0.01 |
| 43 | Mali | 0.33 | 86 | Bolivia | 0.1 | 129 | Bangladesh | <0.01 |

**Appendix 1 Table 6.** List of diseases ordered by the number of deaths caused by each disease during the year 2004 in the world. Data are taken from ***WHO (2008)***. For 8 of these diseases (tagged by a star ^∗^), the number of deaths are estimated using English Wikipedia edition (***Wikipedia, 2018a***).

| Rank | Diseases | Deaths ($\times 10^3$) |
| --- | --- | --- |
| 1 | Pneumonia* | 4000 |
| 2 | HIV/AIDS | 2040 |
| 3 | Tuberculosis | 1464 |
| 4 | Malaria | 889 |
| 5 | Shigellosis* | 700 |
| 6 | Measles | 424 |
| 7 | Foodborne illness* | 420 |
| 8 | Influenza* | 375 |
| 9 | Meningitis | 340 |
| 10 | Salmonellosis* | 268 |
| 11 | Pertussis | 254 |
| 12 | Norovirus* | 200 |
| 13 | Tetanus | 163 |
| 14 | Typhoid fever* | 149 |
| 15 | Hepatitis B | 105 |
| 16 | Syphilis | 99 |
| 17 | Cholera* | 68 |
| 18 | Hepatitis C | 54 |
| 19 | African trypanosomiasis | 52 |
| 20 | Leishmaniasis | 47 |
| 21 | Schistosomiasis | 41 |
| 22 | Dengue fever | 18 |
| 23 | Japanese encephalitis | 11 |
| 24 | Chagas disease | 11 |
| 25 | Chlamydia infection | 9 |
| 26 | Leprosy | 5 |
| 27 | Diphtheria | 5 |
| 28 | Trichuriasis | 2 |
| 29 | Ascariasis | 2 |
| 30 | Poliomyelitis | 1 |
| 31 | Gonorrhea | 1 |

## Footnotes

1 Even if most of the Sepsis are bacterial, it can also be fungal or viral

2 Noted GB in Fig. 6 and Tab. 3.

## References

Al Balushi A, Meeks MW, Hayat G, Kafaie J. Creutzfeldt-Jakob Disease: Analysis of Four Cases. Frontiers in Neurology. 2016; 7:138. https://www.frontiersin.org/artide/10.3389/fneur.2016.00138, doi: 10.3389/fneur.2016.00138.

Armstrong-James D, Meintjes G, Brown GD. A neglected epidemic: fungal infections in HIV/AIDS. Trends in Microbiology. 2014; 22(3):120 – 127. http://www.sciencedirect.com/science/artide/pii/S0966842X1400002X, doi: https://doi.org/10.1016/j.tim.2014.01.001.

Auma MA, Siedner MJ, Nyehangane D, Nalusaji A, Nakaye M, Mwanga-Amumpaire J, Muhindo R, Wilson LA, Boum Y, Moore CC. Malaria is an uncommon cause of adult sepsis in south-western Uganda. Malaria Journal. 2013; 12(1):146. https://doi.org/10.1186/1475-2875-12-146, doi: 10.1186/1475-2875-12-146.

Brin S, Page L. The anatomy of a large-scale hypertextual Web search engine. Computer Networks and ISDN Systems. 1998; 30(1):107 – 117. http://www.sciencedirect.com/science/article/pii/S016975529800110X, doi: https://doi.org/10.1016/S0169-7552(98)00110-X, proceedings of the Seventh International World Wide Web Conference.

Butler D. Publish in Wikipedia or perish. Nature. 2008 dec; https://doi.org/10.1038/news.2008.1312, doi: 10.1038/news.2008.1312.

Callaway E. No rest for the bio-wikis. Nature. 2010 nov; 468(7322):359–360. https://doi.org/10.1038/468359a, doi: 10.1038/468359a.

CDC, Bacterial Meningitis; 2018. https://www.cdc.gov/meningitis/bacterial.html, accessed Sep. 2018.

Chepelianskii AD. Towards physical laws for software architecture. arXiv. 2010; arXiv:1003.5455. http://arxiv.org/abs/1003.5455.

Coquidé C, Lages J, Shepelyansky DL. World influence and interactions of universities from Wikipedia networks. arXiv. 2018 Sep; arXiv:1809.00332. https://arxiv.org/abs/1809.00332.

El Zant S, Frahm KM, Jaffrès-Runser K, Shepelyansky DL. Analysis of world terror networks from the reduced Google matrix of Wikipedia. The European Physical Journal B. 2018 Jan; 91(1):7. https://doi.org/10.1140/epjb/e2017-80570-0, doi: 10.1140/epjb/e2017-80570-0.

El Zant S, Jaffrès-Runser K, Shepelyansky DL. Capturing the influence of geopolitical ties from Wikipedia with reduced Google matrix. PLOS ONE. 2018 08; 13(8):1–31. https://doi.org/10.1371/journal.pone.0201397, doi: 10.1371/journal.pone.0201397.

Encyclopaedia Britannica; 2018. http://www.britannica.com, accessed Aug. 2018.

Eom YH, Aragón P, Laniado D, Kaltenbrunner A, Vigna S, Shepelyansky DL. Interactions of Cultures and Top People of Wikipedia from Ranking of 24 Language Editions. PLOS ONE. 2015 03; 10(3):1–27. https://doi.org/10.1371/journal.pone.0114825, doi: 10.1371/journal.pone.0114825.

Ermann L, Frahm KM, Shepelyansky DL. Google matrix analysis of directed networks. Rev Mod Phys. 2015 Nov; 87:1261–1310. https://link.aps.org/doi/10.1103/RevModPhys.87.1261, doi: 10.1103/RevModPhys.87.1261.

Frahm KM, Shepelyansky DL. Reduced Google matrix. arXiv. 2016 Feb; arXiv:1602.02394. https://arxiv.org/abs/1602.02394.

Frahm KM, Jaffrès-Runser K, Shepelyansky DL. Wikipedia mining of hidden links between political leaders. The European Physical Journal B. 2016 Dec; 89(12):269. https://doi.org/10.1140/epjb/e2016-70526-3, doi: 10.1140/epjb/e2016-70526-3.

Frahm KM, Shepelyansky DL, Wikipedia networks of 24 editions of 2017; 2017. http://www.quantware.ups-tlse.fr/QWLIB/24wiki2017, accessed Jul. 2018.

Giles J. Internet encyclopaedias go head to head. Nature. 2005 Dec; 438:900–901. https://doi.org/10.1038/438900a, doi: 10.1038/438900a.

Karp G, Schlaeffer F, Jotkowitz A, Riesenberg K. Syphilis and HIV co-infection. European Journal of Internal Medicine. 2009; 20(1):9 – 13. http://www.sciencedirect.com/science/article/pii/S0953620508001301, doi: https://doi.org/10.1016/j.ejim.2008.04.002.

Katz G, Rokach L. Wikiometrics: a Wikipedia based ranking system. World Wide Web. 2017 Nov; 20(6):1153–1177. https://doi.org/10.1007/s11280-016-0427-8, doi: 10.1007/s11280-016-0427-8.

Kwan CK, Ernst JD. HIV and Tuberculosis: a Deadly Human Syndemic. Clinical Microbiology Reviews. 2011; 24(2):351–376. https://cmr.asm.org/content/24/2/351, doi: 10.1128/CMR.00042-10.

Lages J, Patt A, Shepelyansky DL. Wikipedia ranking of world universities. The European Physical Journal B. 2016 mar; 89(3). https://doi.org/10.1140/epjb/e2016-60922-0, doi: 10.1140/epjb/e2016-60922-0.

Lages J, Shepelyansky DL, Zinovyev A. Inferring hidden causal relations between pathway members using reduced Google matrix of directed biological networks. PLOS ONE. 2018 01; 13(1):1–28. https://doi.org/10.1371/journal.pone.0190812, doi: 10.1371/journal.pone.0190812.

Langville AN, Meyer CD. Google’s PageRank and Beyond: The Science of Search Engine Rankings. Princeton University Press; 2012.

Leprosy, Syphilis. LEPROSY AND SYPHILIS. The Lancet. 1890; 135(3487):1436. http://www.sciencedirect.com/science/article/pii/S0140673602192002, doi: https://doi.org/10.1016/S0140-6736(02)19200-2, originally published as Volume 1, Issue 3487.

Liberski PP, Ironside JW. Chapter 23 - Prion Diseases. In: Zigmond MJ, Rowland LP, Coyle JT, editors. Neurobiology of Brain Disorders San Diego: Academic Press; 2015.p. 356 – 374. http://www.sciencedirect.com/science/article/pii/B9780123982704000239, doi: https://doi.org/10.1016/B978-0-12-398270-4.00023-9.

Nielsen FÅ. Wikipedia Research and Tools: Review and Comments. SSRN Electronic Journal. 2012; https://doi.org/10.2139/ssrn.2129874, doi: 10.2139/ssrn.2129874.

NIH, Infectious diseases; 2018. https://www.nih.gov/about-nih/what-we-do/nih-turning-discovery-into-health/infectious-diseases, accessed Aug. 2018.

Pappas PG, Lionakis MS, Arendrup MC, Ostrosky-Zeichner L, Kullberg BJ. Invasive candidiasis. Nature Reviews Disease Primers. 2018 May; 4:18026 EP –. http://dx.doi.org/10.1038/nrdp.2018.26, primer.

Pawlowski A, Jansson M, Sköld M, Rottenberg ME, Källenius G. Tuberculosis and HIV Co-Infection. PLOS Pathogens. 2012 02; 8(2):1–7. https://doi.org/10.1371/journal.ppat.1002464, doi: 10.1371/jour-nal.ppat.1002464.

Reagle Jr JM. Good Faith Collaboration: The Culture of Wikipedia (History and Foundations of Information Science). The MIT Press; 2012.

Shane AL. Red Book: 2006 Report of the Committee on Infectious Diseases, 27th Edition, vol. 12; 2006. http://wwwnc.cdc.gov/eid/article/12/12/06-1045, doi: 10.3201/eid1212.061045.

Shannon P, Markiel A, Ozier O, Baliga NS, Wang JT, Ramage D, Amin N, Schwikowski B, Ideker T. Cytoscape: A Software Environment for Integrated Models of Biomolecular Interaction Networks. Genome Research. 2003; 13(11):2498–2504. http://genome.cshlp.org/content/13/11/2498.abstract, doi: 10.1101/gr.1239303.

Sikorska B, Liberski PP. In: Harris JR, editor. Human Prion Diseases: From Kuru to Variant Creutzfeldt-Jakob Disease Dordrecht: Springer Netherlands; 2012. p. 457–496. https://doi.org/10.1007/978-94-007-5416-4_17, doi: 10.1007/978-94-007-5416-4_17.

Syphilis, Leprosy. Syphilis and leprosy. Journal of the American Medical Association. 1899; XXXIII(17):1048–1049. http://dx.doi.org/10.1001/jama.1899.02450690052009, doi: 10.1001/jama.1899.02450690052009.

WHO. The Global Burden of Disease: 2004 Update. World Health Organization; 2008. http://www.who.int/healthinfo/global_burden_disease/2004_report_update/en/.

WHO, Global Health Observatory data repository; 2018. http://apps.who.int/gho/data/node.home.

Wiki4InfectiousDiseases, Wikipedia network of infectious diseases; 2018. http://perso.utinam.cnrs.fr/~lages/datasets/Wiki4InfectiousDiseases/, accessed Sep. (2018).

Wikipedia, List of sovereign states in 2017; 2017. https://en.wikipedia.org/wiki/List_of_sovereign_states_in_2017, accessed Jul. 2018. Wikipedia.

Wikipedia; 2018. https://en.wikipedia.org/wiki/Cholera,\https://en.wikipedia.org/wiki/Typhoid_fever,\https://en.wikipedia.org/wiki/Salmonellosis,\https://en.wikipedia.org/wiki/Shigellosis,\https://en.wikipedia.org/wiki/Foodborne_illness,\https://en.wikipedia.org/wiki/Norovirus,\https://en.wikipedia.org/wiki/Influenza,\https://en.wikipedia.org/wiki/Pneumonia. Wikipedia.

Wikipedia, Immunocompetence; 2018. https://en.wikipedia.org/wiki/Immunocompetence, accessed Sep. 2018. Wikipedia.

Wikipedia, Jenks natural breaks optimization; 2018. https://en.wikipedia.org/wiki/Jenks_natural_breaks_optimization, accessed Jun. 2018. Wikipedia.

Wikipedia, List of infectious diseases; 2018. https://en.wikipedia.org/wiki/List_of_infectious_diseases, accessed Jul. 2018. Wikipedia.

Wikipedia, Mycobacterium; 2018. https://en.wikipedia.org/wiki/Mycobacterium, accessed Sep. 2018. Wikipedia.

Wikipedia, Waterborne diseases; 2018. https://en.wikipedia.org/wiki/Waterbome_diseases, accessed Sep. 2018. Wikipedia.

Zhirov AO, Zhirov OV, Shepelyansky DL. Two-dimensional ranking of Wikipedia articles. The European Physical Journal B. 2010 Oct; 77(4):523–531. https://doi.org/10.1140/epjb/e2010-10500-7, doi: 10.1140/epjb/e2010-10500-7.

